# *Blockbuster* meets the theoretical limit of genome-wide SFS-based inference of recent demography

**DOI:** 10.64898/2026.03.16.712122

**Authors:** Abdelmajid Omarjee, Amaury Lambert, Guillaume Achaz

## Abstract

The molecular diversity of a sample of DNA sequences from the same species is traditionally summarized by the so-called Site Frequency Spectrum (SFS). Past variations in population size leave a trace in the shape of this histogram that several popular programs leverage to infer demography. The ability of inference methods using genome-wide SFS to document recent demography for conservation purposes needs to be evaluated both theoretically and practically. Assuming population size is piecewise constant, we predict analytically that the date of the most recent demographic change that leaves a statistically detectable trace in the SFS is on the order of 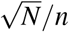 generations, where *n* is the sample size and *N* is the effective population size before the change. We further show that robust parameter estimation is achieved at *N*^3*/*4^*/n* generations for both the date and intensity of the most recent change. We release *Blockbuster*, a deterministic program that reliably infers a demographic scenario with piecewise-constant population size through time from a genome-wide SFS. *Blockbuster* consistently outperforms recent similar programs in accuracy, robustness, and computational time. More importantly, it reaches the theoretical limit on the most recent demographic changes. Finally, we propose a simple yet efficient method to circumvent the presence of population structure, a well-known and pervasive complication that prevents reliable inference of demographic history.

## 2 Introduction

Amid the biodiversity crisis driven by human activities^1^, the IUCN Red List, which assigns conservation statuses to species, remains a cornerstone of conservation biology. Yet, its strong taxonomic biases and its reliance on costly, difficult-to-collect data^2^ highlight the need for automated and scalable approaches capable of evaluating a wider range of organisms and quantifying the status of wild populations^3,4^. The rapid expansion of sequencing technologies^5,6,7,8^ now enables the use of population genetic models and methods^9,10,11,12^ to infer demographic histories directly from genomic data. This influx of cost-efficient sequences has spurred the development of numerous statistical methods aimed at reconstructing effective population size trajectories through time, *N*_*e*_(*t*). Several classes of genomic summary statistics have been proposed to estimate *N*_*e*_(*t*), here estimated from coalescence rates^13^:

1. The first class relies on the distribution of allele frequencies at the genome scale, the Site Frequency Spectrum (SFS), and includes programs such as *Dadi*^14^, *Fastsimcoal2*^15^, *Stairway Plot2*^16,17^ or *FitCoal*^18^. In this article, we introduce a new software called *Blockbuster* which belongs to this category.
2. PSMC-like methods (the second class) leverage sequence variation in coalescence times along the genome^19,20,21,22^.
3. A third class exploits linkage information, including linkage blocks^23,4^ and pairwise covariances between polymorphic sites (*i*.*e*., Linkage Disequilibrium)^24,25,26^.
4. Finally, hybrid approaches combine multiple genomic features^27,28,29,30^.

Although originally designed for reconstructing long-term demographic histories—primarily applied to humans—these approaches hold substantial potential for conservation biology, which focuses on changes unfolding over very recent timescales. Yet, their performance in detecting population size variation in the immediate past (a few to dozens of generations), the window most relevant for conservation assessments, remains insufficiently characterized. Current evidence suggests that different statistics have distinct theoretical limits: PSMC-like (second-class) methods are most informative for ancient periods, whereas LD-based (third-class) approaches are empirically more sensitive to very recent variation^24,31,25^.

Here, we investigate the theoretical limits of SFS-based inferences using tools from mathematical population genetics and test them practically using the *Blockbuster* program. The SFS is a uniquely attractive statistic for demographic inference: it is simple, robust to sequencing errors after minimal filtering (Mirchandani et al., in preparation), rapidly computable from alignments of arbitrary size and avoids the cumbersome need of recombination maps, phasing and/or the presence of long high-quality segments. Although allele frequency distributions have been central to population genetics since its inception^10^, the SFS gained major importance following the derivation of its first moments^32^ and its connection to neutrality tests—including Tajima’s D^33^, Fu and Li’s statistics^34^, and related metrics^35^. It is also at the core of composite-likelihood inference frameworks that can reliably infer model parameters^36,14,15^. Theoretical work has extensively characterized the SFS under coalescent models, including extensions such as changing *N*_*e*_(*t*)^37^, population structure^38^, or multiple mergers^39^. Furthermore, various practical implementations exist to compute SFS from genomic data^40,41,42^. Known limitations, such as identifiability issues, have been documented: an infinite number of distinct demographic histories can produce identical SFS, which impedes identifiability^43^. Although this issue has been challenged in the context of biologically realistic scenarios with effectively infinite genomes and sufficiently large sample sizes, it remains that strikingly different scenarios can be estimated from a single input SFS and major uncertainties in inference across varying population size models decrease only very slowly with more segregating sites^44,45,46^. As a consequence, some inferred ancient demographic events are likely statistical artifacts^18,47,48^. Finally, it is important to stress that small sample size can only help infer coarse-grained size variations, proxied by a small number of epochs^49^. Here, the term *epoch* denotes a time interval on which population size is constant, sometimes also called *block*, hence the name of the program *Blockbuster*.

On the practical side, many existing SFS-based inference programs take hours to days to infer past variations in population size. Furthermore, many rely on Monte Carlo numerical samplers which hamper optimization and reproducibility. To overcome these limitations, *Blockbuster* infers in seconds the best *k*-epoch piecewise-constant demographic model, taking advantage of a deterministic algorithm based on ordinary least squares. *Blockbuster* substantially outperforms most existing programs in both speed and statistical power, reaching the theoretical frontier for detecting recent size changes: 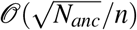 generations, where *N*_*anc*_ is the ancestral effective population size and *n* the sample size, and 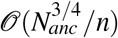 for reliable parameter estimation. We finally show how ignoring high-frequency variants enables robust inference despite pervasive population structure. This simple yet efficient trick addresses a major barrier to the reliable estimation of recent demography.

## 3 Results

We have developed a fully deterministic framework to infer piecewise-constant demography, which we implemented in a new standalone software named *Blockbuster*. This program can be tuned with several options, but here we report results only for the default setting, unless specified otherwise. To assess the performance of the new program, we used real data and a collection of different simulated scenarios, from simple ones (e.g., a single large change in population size over a moderate timescale with a large sample) to challenging ones (small sample size, many epochs or even continuous variations, subtle size changes, and/or the confounding effect of population structure). We also compared its performance with four other popular recent programs that infer past variations in population size from SFS. Finally, we show that *Blockbuster* not only outperforms other programs, but that it actually meets the theoretical limit for recent inference of changes in population size.

### 3.1 Blockbuster method overview and applications

For a given Site Frequency Spectrum (SFS), *Blockbuster* produces identical results across repeated runs. As a result, unlike stochastic approaches such as *Dadi, Stairway Plot2*, or *Fastsimcoal2*, there is no need to perform multiple optimization runs and retain only the best solution. This property is especially advantageous for models with more than three epochs, where stochastic methods typically require extensive computational effort to ensure convergence to a pseudo-optimal solution. Here, we assume a model in which population size *N*_*e*_(*t*) is piecewise constant with (backward) time *t*, also known as a *k*-epoch model, where *k* corresponds to the number of epochs (time intervals of population size constancy). The standard neutral model corresponds to *k* = 1. We leverage the fact that under any *k*-epoch model, the expected SFS can be expressed as a linear combination of the *k* different past sizes (see Supplementary Material Section 2). *Blockbuster* systematically explores all possible solutions from *k* = 1 to a user-defined maximum number of epochs (typically *k*_max_ = 5 or 6). For each value of *k* ∈ {1,…, *k*_max_}, the *k* population sizes corresponding to each epoch can instantaneously be inferred using ordinary least squares, once the *k*− 1 time points of change between epochs are specified. The algorithm thus performs an exhaustive search over a grid of time points, systematically examining all combinations on a relatively coarse grain temporal grid, and then subsequently refines the solution locally from the best candidate identified during this initial global search. For all solutions, a composite likelihood is computed (see Supplementary Material Section 1.1) to select the “best” solution. Likelihood ratio tests (LRTs) are used to statistically select the best *k* as the most parsimonious number of epochs. This approach entirely bypasses all issues related to numerical optimization (assessing convergence from random initial points, heuristic numerical explorations such as gradient descent, etc.) and thus magnifies the advantages of this method.

The overall CPU time can be controlled by the user by setting *k*_*max*_ the maximum value for the number *k* of epochs and the granularity of the time grid. The execution time is mainly determined by the exhaustive search, whose complexity is 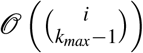, where *i* is the number of points in the time grid and *k*_*max*_ is the maximal number of epochs. In practice, the inference takes from a few seconds to a few minutes on a standard laptop for sample sizes up to *n* = 1, 000.

Several options are implemented in *Blockbuster* allowing it to fit both unfolded and folded SFS. Folded SFSs are typically used when the ancestral state of a segregating site is unknown. One can also truncate the SFS by specifying the index of the last bin to be included. To account for sequencing errors, singletons can be ignored^50^.

As an illustration, we first generated the expected SFS corresponding to a 5-epoch scenario (using Equation 10 of Supplementary Section 2) and used it to infer the demographic history using *Blockbuster*. The inferred history matches the true scenario very closely, demonstrating the power of the approach (Figure 1A).

**Figure 1.**
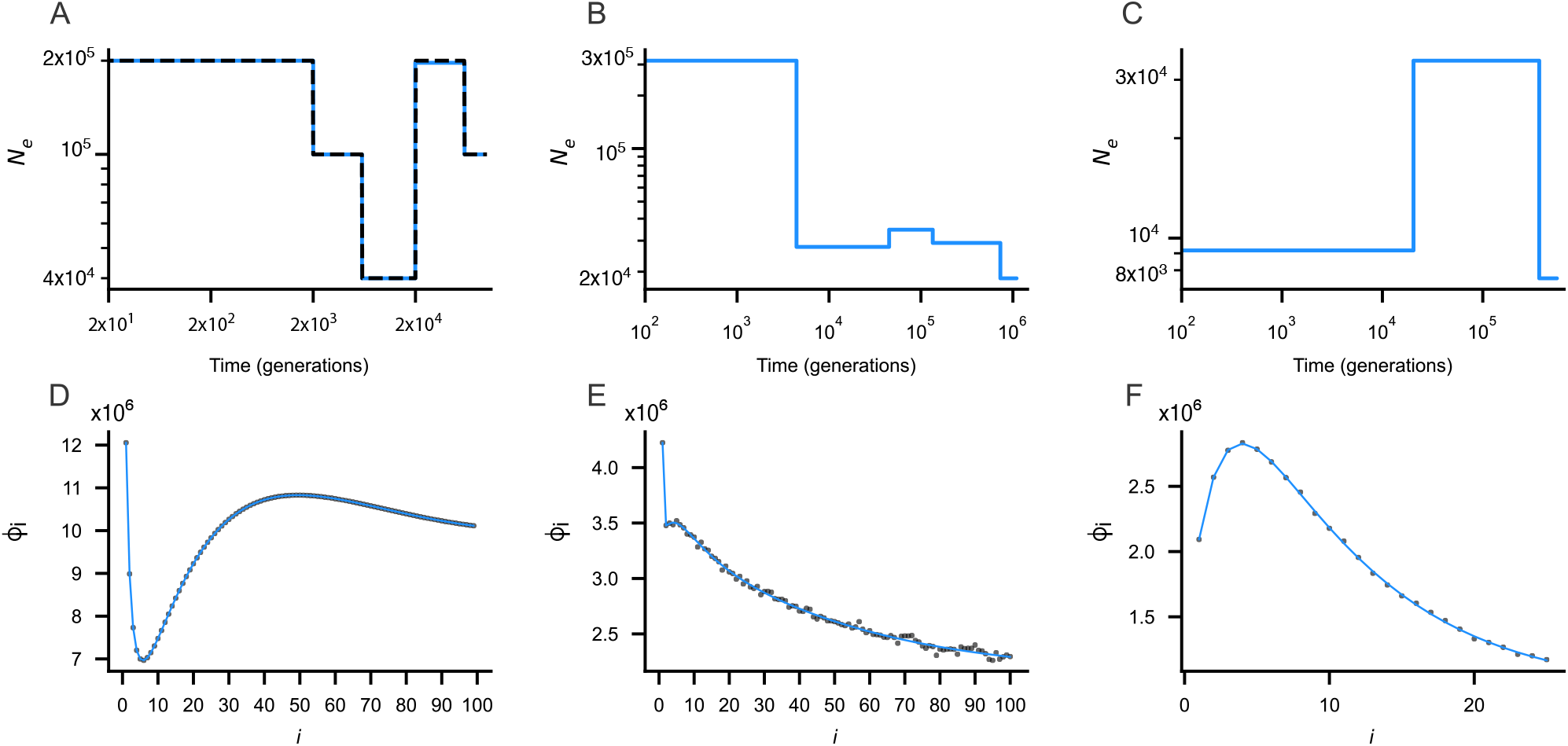
Demographic inference and corresponding transformed SFS (*ϕ*_*i*_ = *iξ*_*i*_) for three datasets. Panels **A– C** show the inferred demographic histories (blue) for: (**A**) a simulated SFS, (**B**) Yoruba, and (**C**) *Gorilla gorilla*. For the simulated SFS with *µL* = 25 (**A**), the true demographic history is shown as a black dashed line for reference. Panels **D–F** show the corresponding transformed SFS: observed (black) and inferred (blue) for the simulated (**D**), Yoruba (**E**), and *Gorilla gorilla* (**F**) datasets.

We also applied the program to real data from a human Yoruba population (2*n* = 208) and to a gorilla population (2*n* = 54), which SFS have already been used in previous studies^45,39^. We truncated them by removing high-frequency variants, that are known to be skewed upward by selection^51,52^, population structure^38,53^, other model misspecification^54,55^, or misorientation errors when the ancestral and derived states of alleles are incorrectly assigned^56^. We kept the variant counts in the 1-100 bins for the Yoruba and in the 1-25 bins for the gorillas. See Methods (Section 5.1) for more details.

The strong excess of singletons observed in Yorubas is captured by a sharp increase in the effective population size in recent times (Figure 1B), which is in accordance with previously published results^57,17,45^. We did not detect any ancient bottlenecks in humans, contrarily to what has been reported elsewhere^18^. The present-day effective size of Yoruba is estimated to be *N*_*e*_(0) ≈ 3 × 10^5^.

For the gorilla subspecies, a 3-epoch demographic scenario is selected, with an initial expansion phase to *N*_*e*_ ≈ 3 × 10^4^ in ancient times followed by a more recent decline down to *N*_*e*_ ≈ 8 × 10^3^, roughly ten thousand years ago (Figure 1C). We infer no trace of very recent decline, likely due to the small sample size (see Section 3.3), even when considering scenarios with *k* = 4 or 5 epochs (data not shown).

### 3.2 Benchmark analysis

We compared the power and limitations of *Blockbuster* to those of four other popular programs: *Dadi*^14^, *Fastsimcoal2*^15^, *Stairway Plot2*^16,17^ and *FitCoal*^18^.

#### 3.2.1 CPU time and Memory

We evaluated the CPU time and memory usage of all programs (see Methods section 5.2 for details). As expected, while fitting the 5-epoch model of Figure 1A, the CPU time of *Blockbuster* increases approximately linearly with sample size (Figure 2A). It takes 10 seconds for *n* = 20 haploid genomes and about 100 seconds for *n* = 200. *Dadi* is slower by approximately one order of magnitude, whereas *Stairway Plot2* and *Fastsimcoal2* are two orders of magnitude slower. Although the execution time of *FitCoal* is similar to that of *Dadi*, we infer models with up to *k*_*max*_ = 5 epochs for the former, compared to *k*_*max*_ = 3 for the latter (even though the true scenario has 5 epochs). An important feature of *Blockbuster* is that its execution time does not depend on numerical convergence, whereas other programs can be more or less time-consuming depending on the underlying model and the smoothness of the associated likelihood surface. In terms of memory usage, consumption increases linearly with sample size for *Blockbuster* and *Stairway Plot2*, but remains approximately constant for the other three programs. For these latter programs, the most memory-intensive steps do not depend on sample size, but likely on external libraries. While *Stairway Plot2* and *FitCoal* require more than 1 Gb of memory regardless of sample size, the three other programs require between tens to hundreds Mb (Fig. 2B), which is now possible on standard laptops. Therefore, *Blockbuster* appears to offer an excellent compromise between model complexity, time consumption, and memory usage.

**Figure 2.**
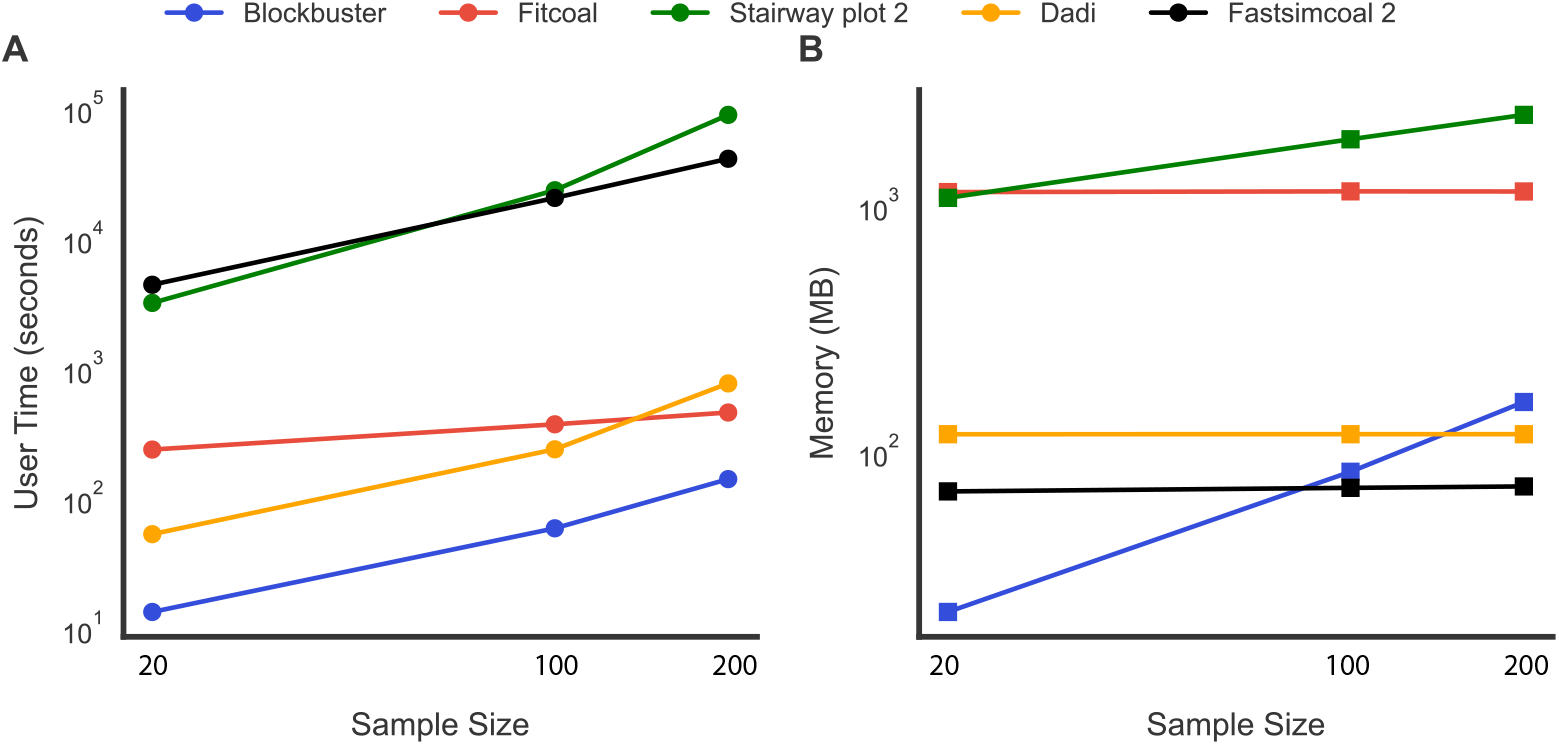
Computational efficiency and memory requirements for 5-epoch demographic models. Panels **A, B** compare runtime and memory usage across five demographic inference programs (*Blockbuster, FitCoal, Dadi, Stairway Plot2*, and *Fastsimcoal2*) for inferred 5-epoch histories at sample sizes of *n* = 20, 100, 200 haplotypes.

Importantly, execution time of *Blockbuster* is *not* linear with respect to the temporal grid size or the number of epochs. For 5-epoch models, halving the temporal grid size (from 100 to 50) results in an approximately ten-fold reduction. In contrast, increasing the number of epochs from 5 to 7 (for a grid of 50 time points) results in an approximately fifty-fold increase in execution time (Figure S1A). Consequently, reducing the grid size can be advisable when increasing the maximum number of epochs. *Blockbuster* nonetheless remains faster than the other programs in all tested configurations. Finally, memory usage of *Blockbuster* does not depend on the number of parameters inferred (compare the 5- and 7-epoch cases in Figure S1B), but increases linearly with the temporal grid size.

#### 3.2.2 Statistical power

To evaluate the performance of *Blockbuster*, we conducted intensive simulations using *k*-epoch models with *k* = 2, 3, 4, or 5 epochs of various timings and intensities. We even explored scenarios with continuous linear or exponential changes in *N*_*e*_ that, by design, cannot be properly inferred as such by *Blockbuster*.

For each demographic model and each sample size, we generated the exact expected SFS (*E*[*ξ*_*i*_], see Supplementary Material, Section 2 Eq 10) and used it as input for the five programs. To complement this, we simulated 40 additional “noisy” SFS, populating each bin of the SFS (*ξ*_*i*_) with a Poisson random variable whose mean (and thus variance) was set equal to the expected value (*E*[*ξ*_*i*_]). This was motivated by our desire to assess the impact of noise on the quality of inferences. Accordingly, all five programs were also evaluated on noisy SFS. We acknowledge that the added noise is not necessarily realistic, as it does not account for recombination, which generates correlated genealogies along the genome^58^, but it still provides information on the robustness of inferences.

For each input SFS, we scored each demographic inference by the *L*^2^ distance between true demography and inferred demography (Root Mean Square Error, or RMSE). We ranked the five programs from the most accurate (smallest RMSE) to the least accurate (largest RMSE).

For the 2-epoch models, we tested three sample sizes (*n* ∈ {20, 100, 500}), four change times (*T* ∈ {0.001, 0.01, 0.1, 1} × *N*_pres_ generations), and six different ancestral sizes (*N*_anc_ = *ρ* × *N*_*e*_(0), with *ρ* ∈ {0.1, 0.5, 0.9, 1.1, 5, 10}). In total, we computed 3 × 4 × 6 = 72 distinct scenarios (parameter combinations) for *k* = 2 and, consequently, 72 × 41 = 2,952 input SFS, where 41 is the number of replicates (1 expected SFS plus 40 noisy SFS). We also studied eleven 3-epoch models, two 4-epoch models and three 5-epoch models, with 40 + 1 replicates each. See Methods (section 5.3) for details.

We describe results when the input dataset is the expected SFS. Results for the seventy two 2-epoch scenarios (Table 1) show that a version of *Blockbuster* using likelihood penalization (see Supplementary Material, Section 3.4) overall outperforms the other programs, and that *FitCoal* also displays very good performance. Results for the 3-, 4- and 5-epoch scenarios (Table 1) show an even higher relative performance by *Blockbuster*. For 5-epoch models *Stairway Plot2* also displays a good RMSE-based score.

**Table 1:**
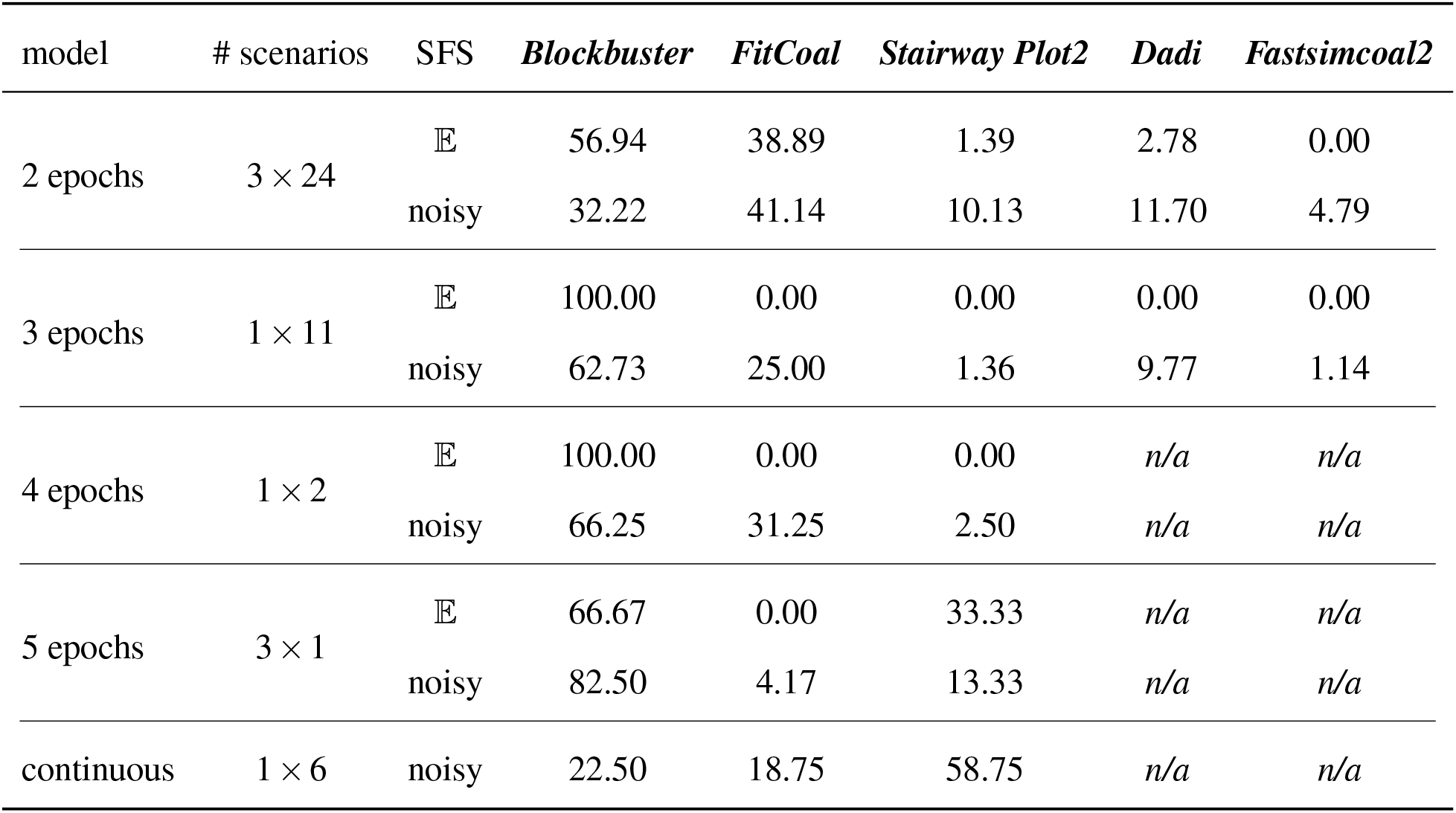
Benchmark. Percentage of inferences for which each program is ranked first, according to the Root Mean Square Error (RMSE). Results are summarized by class of models (*k*-epochs or continuous). The reported number of scenarios per class is the product of the number of sample sizes × the number of different intensities. For each type of model, results for the expected SFS (𝔼) are shown on top, and results for “noisy” SFS (40 replicates per scenario) are shown on the bottom. For partly continuous models, only noisy replicates are represented. Due to long execution times, *Dadi* and *Fastsimcoal2* were only run for 2- and 3-epoch models. Here *Blockbuster* uses likelihood penalization.

| model | # scenarios | SFS | *Blockbuster* | *FitCoal* | *Stairway Plot2* | *Dadi* | *Fastsimcoal2* |
| --- | --- | --- | --- | --- | --- | --- | --- |
| 2 epochs | $3 \times 24$ | $\mathbb{E}$ | 56.94 | 38.89 | 1.39 | 2.78 | 0.00 |
|  |  | noisy | 32.22 | 41.14 | 10.13 | 11.70 | 4.79 |
| 3 epochs | $1 \times 11$ | $\mathbb{E}$ | 100.00 | 0.00 | 0.00 | 0.00 | 0.00 |
|  |  | noisy | 62.73 | 25.00 | 1.36 | 9.77 | 1.14 |
| 4 epochs | $1 \times 2$ | $\mathbb{E}$ | 100.00 | 0.00 | 0.00 | *n/a* | *n/a* |
|  |  | noisy | 66.25 | 31.25 | 2.50 | *n/a* | *n/a* |
| 5 epochs | $3 \times 1$ | $\mathbb{E}$ | 66.67 | 0.00 | 33.33 | *n/a* | *n/a* |
|  |  | noisy | 82.50 | 4.17 | 13.33 | *n/a* | *n/a* |
| continuous | $1 \times 6$ | noisy | 22.50 | 18.75 | 58.75 | *n/a* | *n/a* |

When sampling noise is added, *Blockbuster* performs better than the other programs for 3-to 5-epoch models. For 2-epoch scenarios, *FitCoal* performs better than *Blockbuster* with or without penalization. We conjecture that that this relates to two features of *FitCoal*. First, *FitCoal* constrains the maximum relative variation between consecutive blocks to a tenfold change and second, it has for 2-epochs, an excellent optimization procedure. However, as the number of parameters to be inferred increases, the advantage of the exhaustive (yet discrete) search implemented in *Blockbuster* becomes conspicuous.

Results with the unpenalized likelihood function for *Blockbuster* (Table S1) show a slightly different pattern. In general, penalization tends to bias the inference when the population size change is very recent relative to the sample size. Since likelihood penalization improves performance in most piecewise-constant scenarios (by preventing overfitting), we kept this option as default.

For scenarios with continuous variations (Table 1 and Table S4), *Blockbuster* and *FitCoal* have larger RMSE than *Stairway Plot2*. Indeed, the output of the latter better mimics continuous variations thanks to its *post-hoc* subsampling-plus-smoothing procedure. On a side note, *Blockbuster* also includes a sampling option and its relevance along with other strategies to fit complex models (e.g., zigzag model) is discussed in the Supplementary Material (Section 5 and Figure E1).

Overall, the benchmark shows that the order of performance is: 1) *Blockbuster*, 2) *FitCoal*^16,17^, 3) *Stairway Plot2*^18^, 4) *Dadi*^14^, and 5) *Fastsimcoal2*^15^.

Although handy, RMSE does not specifically capture the presence of spurious size variations that some programs may infer. We therefore complemented the RMSE analysis with visual representations of the inferred demographies, reported in Extended data (see Figures S3, S4, S5 and S6). The three leading programs (*Blockbuster, Stairway Plot2*, and *FitCoal*) have their respective advantages and disadvantages. For instance, *Stairway Plot2* tends to overestimate the number of epochs and shows little robustness for recent demographic changes. *FitCoal* encounters difficulties when fitting multi-epoch models and infers spurious ancient bottlenecks (see Figures S2, S3, S4, and S5). Finally, unpenalized *Blockbuster* can exhibit high variance across replicates (Figure S7), whilst penalization can lead to slight underestimation of size variations (Figure S8).

### 3.3 At the frontier of recent times

After comparing *Blockbuster* with other programs, we explored its performance in a 2-epoch model where the change is very recent, a scenario that is especially relevant to conservation concerns. In this aim, we studied theoretically how recent a change can be and still be statistically detected from the SFS. We were particularly interested in the dependence of this theoretical limit on sample size, population size and intensity of population size variation.

By theoretical considerations combining mathematical population genetics and probability theory (see Supplementary Material Section 4), we computed the expected absolute difference at each bin between an expected SFS corresponding to a 2-epoch scenario and a 1-epoch standard neutral SFS (Proposition 4.1 in Supplementary Material). The sum over all bins of these differences (*L*^1^-distance between the two expected SFS) can be expressed as:

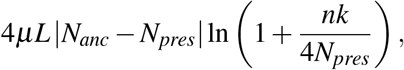

where *n* is the haploid sample size, *N*_*pres*_ the present effective size and *N*_*anc*_ the ancestral effective size before the change, that occurred *k* generations ago. Parameters *µ* and *L* stand for the mutation rate per site per generation and the genome length, respectively. We then showed that a demographic change affecting only the relative abundance of singletons is sufficient to reject the null model provided this change is older than 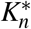 generations (Proposition 4.4 in Supplementary Material), with

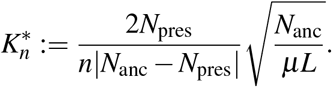

From this equation, all other things being equal, we can, for instance, conclude that a population *decline* of 33% shares the same limit of detection as a population *growth* of 100%.

We further showed that parameters can only be reliably identified if the demographic change is also affecting the doubleton bin. Mathematically, this translates into a demographic change that is older than 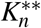 generations (Proposition 4.9 in Supplementary Material), with

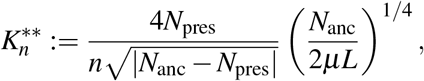

Between these two limits, the inferences are characterized by a very high variance into the parameter values estimated across replicates (Figure S7), despite a correct inference of the direction (*i*.*e*. growth *versus* decline).

From these derivations, we conclude that, once sampling noise is added, “reasonable” changes (*N*_*pres*_ = *c*× *N*_*anc*_, where *c* is fixed) become significant when the change occurs on a time that is at least 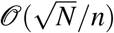 generations ago. We note the proportionality to 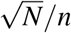, and not to *N/n*^2^, which corresponds to the timescale of the most recent coalescence events. Similarly the second limit, which is the most recent time needed to identify the parameters, is on the order of 𝒪 (*N*^3*/*4^*/n*). Assuming that all other terms in the equality are of order one, this implies that for an ancestral effective population size of *N*_*anc*_ = 10^4^ and a sample size of *n* = 10, the number of generations over which a change can be reliably detected is on the order of *k* ≈ 10 generations for 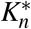 and good parameter estimation of *k* ≈ 100 generations for 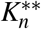. A sample size of *n* = 100 translates into *k* ≈ 1 generations for 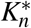 and *k* ≈ 10 generations for robust parameter estimation. Those values, surprisingly small at first sight, are highly relevant for conservation biology.

We obtained further derivations yielding an explicit analytical expression for the power (fraction of true negatives) of the LRT (Proposition 4.5 in Supplementary Material), shown in Figure 3 as a function of the time at which the change occurs.

**Figure 3.**
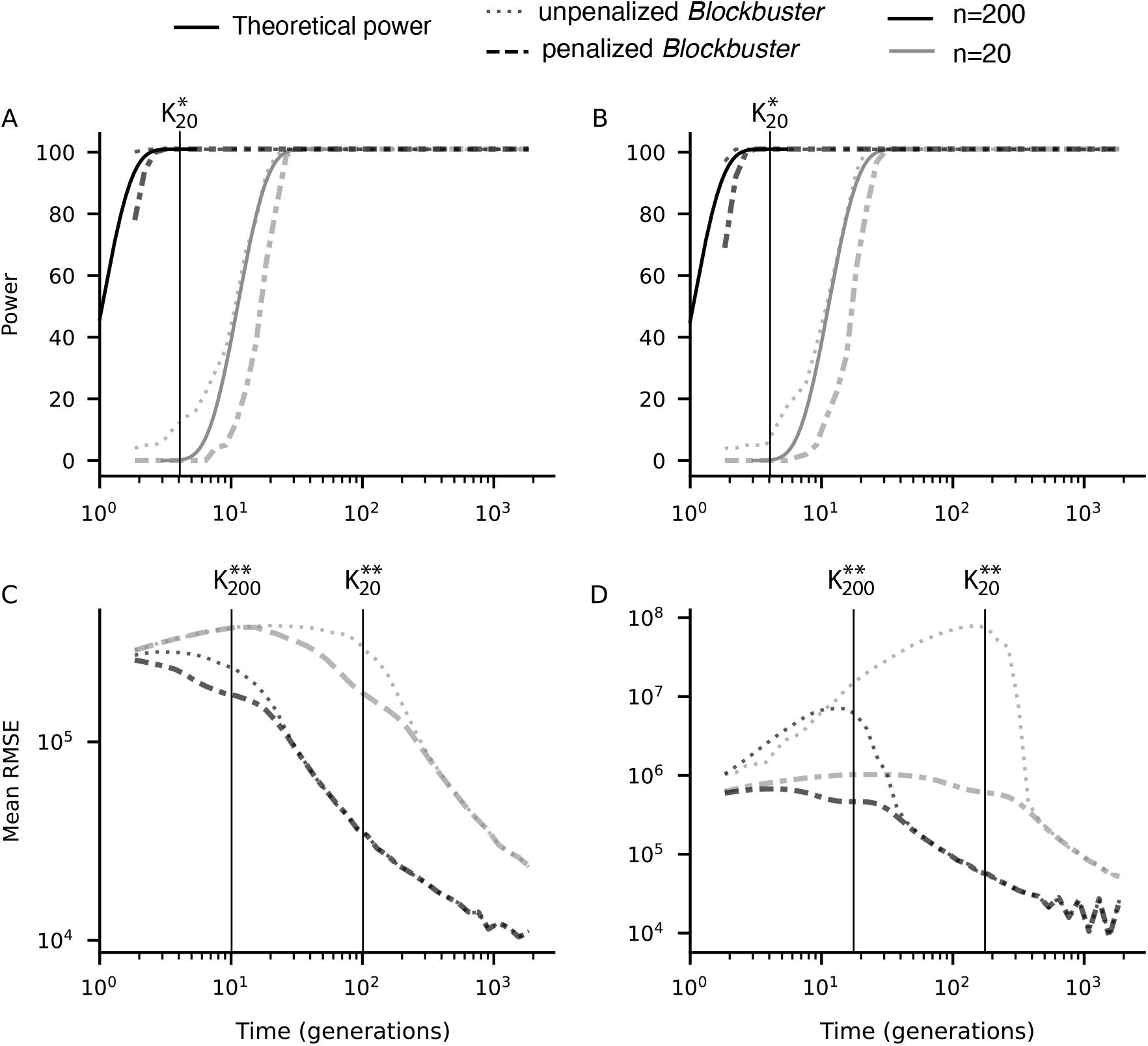
Power and inference accuracy over time for 2-epoch models as a function of the time of change. Left panels (A, C) correspond to two-epoch models with a population decline, whereas right panels (B, D) show models with a population expansion. Power is defined as the proportion (out of 100 replicates) for which the null model is rejected. Continuous curves in panels A and B represent the theoretical power curves, whose expression depends on 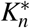, while dotted lines correspond to *Blockbuster* with penalized (dashed lines) or unpenalized (dotted lines) likelihood. The mean RMSE quantifies the reliability of parameter estimation when compared to the true demographic trajectories. Vertical lines represent 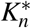 and 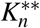 (in panels A and B, 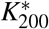 = 0.39 is not represented). Results are shown for sample sizes *n* = 20 (grey) and *n* = 200 (black).

To illustrate these mathematical predictions, we simulated 2-epoch models with different times of change ranging between 1.85 and 1850 generations, for both a 100% population growth and a 33% population decline with the same ancestral size (*N*_*anc*_ = 15,000, *µ* = 1.3 × 10^−8^ an *L* = 3 × 10^9^). For each model, we performed 100 noisy replicates. We then measured a) the fraction of replicates where the null model was rejected and b) the mean RMSE. Inferences were performed with or without likelihood penalization. Results are shown in Figure 3.

First, we note that the empirical power of the *Blockbuster* program meets the mathematical theoretical limits. Second, as predicted, the power curves are almost identical for a population decline of 100% and a population expansion of 33% (Figure 3A, C). Importantly, the time at which it becomes possible to reject the null-model can be extremely recent and clearly in the range relevant for conservation purpose: for a sample size of 20, the theoretical and empirical powers are good (> 0.8) at 20 generations in the past, and for a sample size of 200, power is already maximal at 1 generation in the past. Third, the RMSE curves (as a proxy of the reliability of parameter estimation) show an illuminating pattern. Without penalization, RMSE tends to perform a maximum at intermediate values of the time of change, which points to a large variance in the inferred sizes. This is caused by an identifiability problem (similar statistical signal for a recent, large change as for a distant, moderate change, as illustrated in Figure S7), which inflates the average RMSE. Interestingly the use of penalization almost completely removes this effect (Figure S8), leading to decreasing RMSE with change time. Penalizing the likelihood results in a slight loss of power to detect the change (Figure 3A, B), but in a substantial reduction of RMSE when times are on the order of 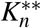. RMSE for more distant past changes do not differ with and without penalization.

Together with Figure S7, these curves confirm the inverse proportionality between time of detectability and sample size: the power curves are shifted by a tenfold factor between *n* = 20 and *n* = 200 haploid genomes. Both the statistical power and the RMSE curves closely match the theoretically identified thresh-olds.

### 3.4 Inferring size variation despite structure

Finally we address a common problem that impedes inference of size variation: the confounding effect of population structure^59,60,13,61^. Through the analysis of simulated scenarios with both population structure and past variations of population size, we came to the conclusion that in many cases, simply ignoring the high frequency bins of the unfolded SFS during the inference, results in correct estimations of the population sizes.

We simulated two standard structured demographic models that combine past variations of population size with population structure. The exact scenarios, inspired from previous studies^62,63,64^, were developed within the *stdpopsim* library^65,66^.

The first scenario, illustrated in Figure 4, corresponds to the prefered scenario for the two orangutan subspecies of Sumatra (*Pongo abelii*) and Borneo (*Pongo pygmaeus*)^62^. *P. abelii* is thought to have diverged from *P. pygmaeus* about 20, 000 generations ago. The past isolation was then followed by asymmetric gene flow, with a pygmaeus-to-abelii bias of approximately twofold (migration rates of 1.10 × 10^−5^ per generation from *P. pygmaeus* to *P. abelii*, and 0.67 × 10^−5^ in the opposite direction). For each simulation, an SFS of 2*n* = 100 haploid genomes were generated for each subspecies using *msprime*^67^ that generates an ARG and hence mimics the natural noise due to recombination (see Methods Section 5.4).

**Figure 4.**
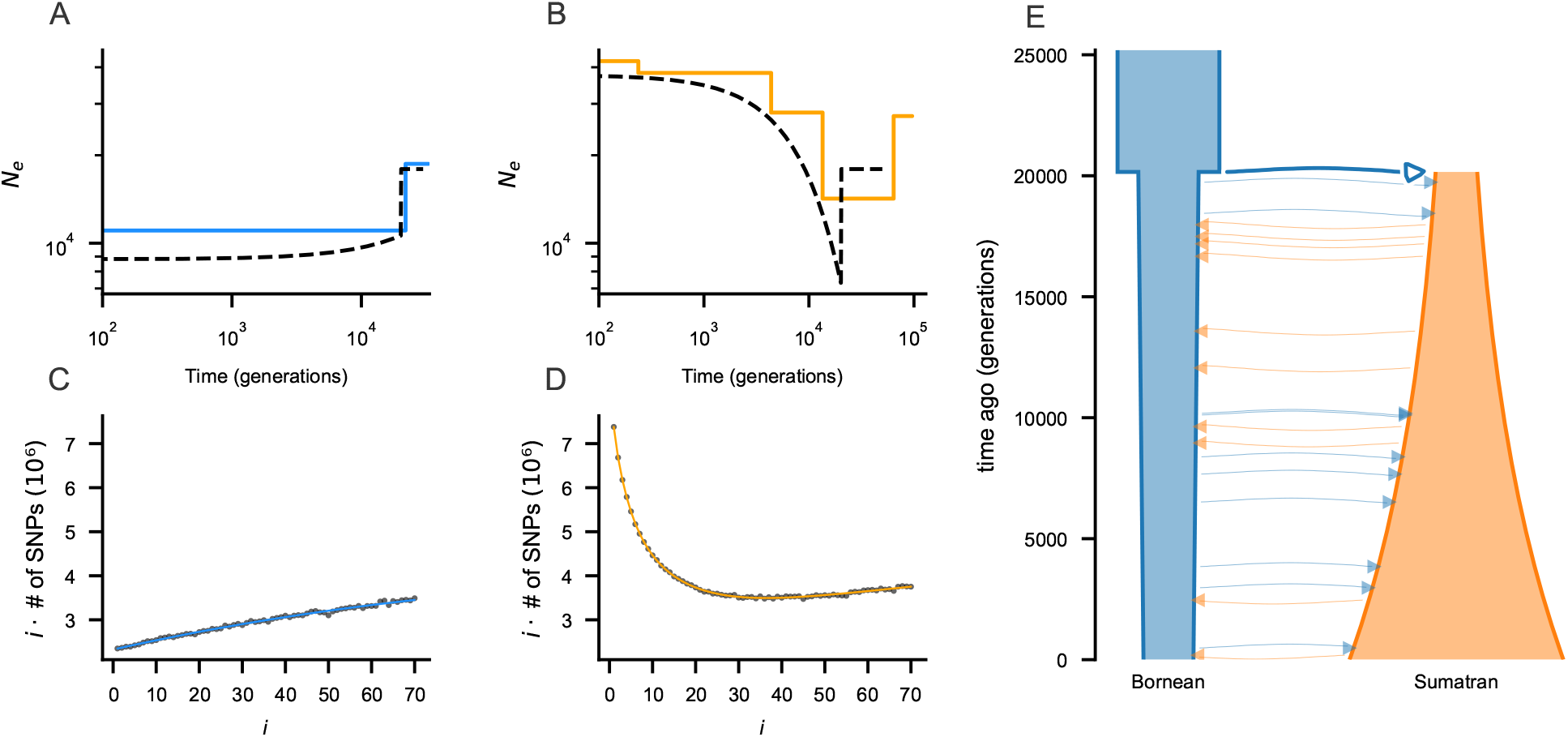
Demographic inference from simulated Site Frequency Spectra (SFS) generated with the *stdpopsim* library for orangutans, while ignoring variants which frequencies exceed 70*/*100. Panels **A–B** show the demographic histories inferred by *Blockbuster* (colored) from a single simulated SFS of the Bornean (**A**) and Sumatran (**B**) populations; the black dashed curves correspond to the true simulated scenarios. Panels **C–D** display the corresponding transformed truncated SFS (black dots) and the fitted SFS infered by *Blockbuster* (colored line). Panel **E** displays a graphical representation of the scenario.

As reported in the literature^38^, isolation with migration scenarios result in an excess of high-frequency variants which obscure the inference of variations of population sizes (Figure S9). Consequently, a simple solution is to ignore the high-frequency bins of the SFS during the inference. Indeed, *Blockbuster* approximately recovers the changes of *N*_*e*_ while only considering the variants which frequencies is equal or lower than 70*/*100 (Figure 4).

We complemented this analysis by simulations where migration rates were null immediately after the split (Figure S10) (isolation, no migration). In this case, we fully recover the correct pattern of past variations using the complete SFS. In comparison, the inference based on a truncated SFS when migration is ongoing is slightly less accurate, suggesting that although the demography trends are mostly well inferred, structure still alters the parameter estimation. We conclude that although truncated SFS can be used to robustly estimate the general trends of *N*_*e*_ variations, the exact parameter estimations should be considered with more caution.

We further confirmed that the use of unfolded truncated SFS enables the recovery of the correct trends of variation of population size using a scenario describing the out-of-Africa model of humans^63,64^ (Figures S12, S11 and S13).

## 4 Discussion

In this article, we characterize the theoretical and practical limits of genome-wide SFS to infer the past variations of population size *N*_*e*_(*t*) though time *t* from a sample of *n* haploid genomes. We introduce *Blockbuster*, a fast, robust, and flexible program designed to infer a piecewise-constant *k*-epoch model from arbitrarily large site frequency spectra with minimal prior specification.

Through an extensive simulation-based benchmark, we demonstrate that, in contrast to other commonly used programs that take SFS as input, *Blockbuster* systematically converges to a unique and accurate demographic scenario, in the form of a series of *k* time intervals or epochs, each supporting a distinct population size. This desirable property is achieved because *Blockbuster* does not rely on Monte Carlo sampling to compute expected SFS nor gradient-based numerical optimization to fit parameters. We subsequently show that when the SFS provided as input is the exact expected SFS, for samples of reasonable size, *Blockbuster* outputs a demographic scenario which is *always* consistent with the true one (correct number of time changes, direction and intensity of changes). This ensures that the deviation to the true scenario, of the scenario inferred from noisy SFS, is mainly due to sampling noise of sites and not to any systematic algorithmic bias. This reassuring stability property is not observed for the other benchmarked programs. In addition, we observed that *Stairway Plot2* tends to infer incorrect, recent size variations, and that *FitCoal* tends to infer incorrect ancient bottlenecks (a trend reported and discussed elsewhere^18,47,48^).

Using coalescent theory, we characterize mathematically how recent can be the change in a 2-epoch model to be detected from the SFS, depending on sample size and on population sizes. We show that *Blockbuster* meets the theoretical limit of 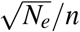 generations, a limit that is highly relevant to conservation biology (1 generation for a sample of *n* = 100 for a species with *N*_*e*_ = 10^4^).

However, when only singletons are statistically different, a classical problem of identifiability arises: a recent but intense change in size is equivalent to a less recent more moderate change. When doubletons are also statistically different, which happens for a change time of at least 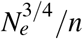 generations, the respective effects of intensity and time decouple, enabling reliable parameter estimation. On a side note, this also suggests that opting out singletons^50^ will shift the limit of detection from 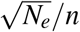 generations to 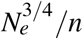 generations, which quantifies the loss of power at detecting recent changes caused by sequencing errors.

Observe that both of the time limits (detection and identifiability) are proportional to *N*^*b*^*/n* where *b* is smaller than 1. This may explain the incorrect recent variations in population size obtained with *Stairway Plot2*, as the first time point used by this program corresponds to the first coalescence time, which is on the order of 1*/n*^2^. In practical terms, this suggests that, in a conservation context, it is possible to detect changes of interest over the scale of a few generations. It obviously becomes easier under certain conditions: an intense change of population size, a large sample size, and a small absolute effective population size. This result has nonetheless to be taken with care as we consider a model with one single change of population size. When inferring more complex demographic histories, quantitative requirements on sample size do not only relate to the date of the most recent change to infer, but also to the number of demographic epochs to infer^49^.

Using similar theoretical arguments, it would be interesting to investigate the limits of other statistics used in population genetic inference, such as linkage disequilibrium^24,25,26^ or the distribution of coalescence time distributions along the genome^19,20,21,22^. Characterizing the theoretical limits of several different genomic metrics could help developing a joint statistical framework, possibly using machine learning, combining them to push the limits of demographic inference beyond the one presented here, using solely the SFS^31^.

We finally show that ignoring high-frequency variants in the inference procedure (*Blockbuster* has an option to simply ignore bins of the SFS) enables us to infer changes of population size despite the presence of population structure. As an example, we illustrate how to use *Blockbuster* (and other SFS-based inferences) despite the presence of ongoing migration between subpopulations. Using the unfolded truncated SFS of each subpopulation, we can recover their general demographic trends, despite a mild bias in the estimation of parameters (intensity and timing of changes) caused by migration. Partitioning a dataset into subpopulations is easily achievable using standard analyses such as hierarchical clustering or PCA. Once the SFS is generated for each subpopulation, a potential hint of ongoing migration is an over-representation of high frequency variants, easily spotted as a in the transformed SFS (*ϕ*_*i*_ = *iξ*_*i*_). In this case, we recommend a simple rule of thumb which is to ignore the last third of the SFS in the inference.

However, one should carefully interpret such inferences in light of naturalistic knowledge about the population of interest. First, complex spatial structures, such as in island or stepping-stone metapopulation models, can have more pervasive confounding effects that are much harder to disentangle depending on the sampling strategy^13^. Second, *U*-shape (excess of low and high frequency variants) can arise in other contexts than population structure, in particular as an effect of linked selection resulting from hitchhiking on recurrent selective sweeps^68^, a regime that has been coined *genetic draft*^69^. The underlying coalescent process is not anymore well approximated by a standard coalescent with varying population size but more so by a Multiple Merger Coalescent (MMC)^54^. Using simulations of beta-coalescent, we show that even when the high frequency bins are ignored, MMC still inflates the relative abundance of low frequency variants that is then interpreted as a signal of growing population by *Blockbuster* (Figure S14). This erroneous inference of population growth will be apparent in all popular methods (among which the ones tested here), that all rely on a binary coalescent.

Several functionalities will be added to future versions of *Blockbuster*. First, a robust subsampling or bootstrapping strategy or a Bayesian mode could be implemented to fit better continuous and realistic demographic scenarios, similarly as *Stairway Plot2* does. In this line, a bootstrapping method using ARG simulations has been proposed to better take into account biological processes such as recombination and varying ancestries along the genome (Loay et al., in preparation). Second, we will enrich *Blockbuster* to use serial samples with one SFS per sampled time point. The main challenge then becomes to express sampling times in units of 2*N*_*e*_(0) generations to be directly implemented within the *Blockbuster* framework. The prior knowledge of the current population sizes needed to rescale time suggests that an EM-like algorithm would be adequate. Finally, we could include a high-precision computation libraries to handle unusually large sample sizes (*n* > 1, 000).

To conclude, we have been characterizing the power of genome wide SFS-based to infer recent variation of population size and develop a stand-alone computer program that meets these theoretical limits, based on classical statistics (ordinary least squares) and champions CPU time (thus keeping the carbon footprint to lower cost). Remains to catalog and analyze many genome-wide SFS of real biological species to assess the empirical robustness of the strategy for conservation purposes.

## 5 Method

### 5.1 Real data application (humans and gorillas)

To scale effective population sizes and times, we applied the following parameters: a mutation rate of *µ* = 1.2 × 10^−8^ per site per generation^70,71^, a genome length of *L* = 2.46 gigabases for humans and 2.6 gigabases for gorillas, and a generation time of 30 years for humans^72^ and 19 years for gorillas^71^. The human genome length was rescaled to account for both inaccessible regions of the genome and misoriented SNPs^45^. Because this information was not available for Gorillas, we used 85% of the genome length value as for the Yoruba case.

### 5.2 CPU time and Memory

First, to evaluate the efficiency of the programs in terms of both CPU time and memory usage, we applied the five programs to the 5-epoch model shown in Figure 1A, with three different haploid sample sizes: *n* ∈ {20, 100, 200}. The results shown in Fig. 2 were obtained using the recommended hyperparameters for *Stairway Plot2* (200 bootstrap replicates and three different numbers of sampled breakpoints), *Fastsimcoal2* (40 optimization cycles in the EM algorithm and 1, 000, 000 samplings in the Monte Carlo simulation steps), and *Dadi* (100 numerical optimizations from random starting points). For *FitCoal*, we fitted piecewise-constant models without any exponential phases. For *Blockbuster*, we used a grid size of 100 for the coarse-grained search and *k*_*max*_ = 5. Note that the reported memory corresponds to the maximum memory usage over the entire execution, and that the reported times for *Dadi* and *Fastsimcoal2* are the sums of the inference times for the 2- and 3-epoch models. For *FitCoal*, the reported times correspond to models ranging from 1 to 5 epochs.

### 5.3 Statistical Power

#### 5.3.1 Input SFS

For all *k*-epoch models, we computed the exact expected SFS (using Equation 10 in Section 2, Supplementary Material) and complemented it with 40 “noisy” SFS, using a Poisson random variable for each bin, with a mean equal to *E*[*ξ*_*i*_] of Equation 10. On the contrary, the models with an initial linear or exponential growth phase followed by a plateau were simulated using a home-made simulator implementing the Monte Carlo method of generating the SFS from a given trajectory, as described in^37^. For all models, except the 5-epoch model, we used the same initial value of *θ* (0), set to 10^6^. For the 5-epoch model, the initial value of *θ* (0) was set to 2 × 10^7^.

#### 5.3.2 Program options and arguments

To perform a large number of demographic inferences within a reasonable time frame (*i*.*e*., days), the options used in all programs do not follow the recommended settings, for models with 2 or 3-epochs for *Fastsimcoal2, Dadi*, and *Stairway Plot2*. Specifically, we performed 30 independent optimizations for *Dadi*, 20 replicates for *Stairway Plot2*, and 100,000 samples for *Fastsimcoal2*. For models with more than three epochs or including a continuous component, 200 replicates were used for *Stairway Plot2*. Furthermore, when parameter ranges were required, they were kept as broad as possible.

For *FitCoal* and *Blockbuster* including a penalization term in the likelihood function no minimum or maximum bounds were specified for times; therefore, the default settings were used. For the unpenalized *Blockbuster*, the minimum time was set to 5 × 10^−4^*N*_*pres*_ generations. For *Dadi*, the relative change between two epochs was fixed to a ten-fold difference, whereas the duration of a given epoch was constrained to lie between 10^−4^ and 10*N*_*anc*_ generations. Finally, for *Fastsimcoal2*, parameter ranges expressed in numbers of generations and effective population sizes are required. Effective population sizes were therefore set between 100 and 100,000, while times were set between 5 and 10,000 generations. Note that for this tool, the upper bounds are used only for the initial random sampling, and larger values can still be inferred. The SFS was scaled using a *µL* value of 12.5.

To quantify the performance of the programs beyond visual inspection, we computed an RMSE (Root Mean Square Error) as a measure of distance between the true simulated demography and the demography inferred by the programs. The RMSE was computed on a grid of 1,000 time points on a logarithmic scale (uniformly spaced from *t* = 0.0001 to *t* = 2). The smaller the RMSE, the closer the inferred demography to the true one. To compute the distance metric between the simulated model and the inference results, values were expressed in terms of population mutation rates and times in units of 2*N*_*e*_(0) generations in the simulated scenario. This ensures that the relative positions of the inferred and simulated models correspond correctly when times are expressed in generations.

### 5.4 Structure and stdpopsim

*stdpopsim* provides preset parameters corresponding to plausible demographic scenarios. We simulated the models using the *msprime* engine. For the orangutan, the genome size was set to *L* = 2.67 × 10^9^, divided into 22 autosomes with their real sizes, and a mutation rate *µ* = 2 × 10^−8^ per site per generation. For humans, the genome size was set to *L* = 2.88 × 10^9^, divided into 23 autosomes with their real sizes, and a mutation rate *µ* = 2.36 × 10^−8^ per site per generation. Simulations were performed using the recombination rates provided by *stdpopsim* for these species, with one constant recombination rate per chromosome.

## Acknowledgments

The authors thank Arno Siri-Jegousse for valuable insights into the mathematical development; Elise Kerdoncuff for providing the Yoruba and Gorilla SFS data; and Pol Mourleix and Anis Toussirt for their contributions to the benchmark framework. We also thank Hossameldin Loay and Maël Legouellec as early users for their feedbacks on the *Blockbuster* program. AO and GA thank the Fédération Nationale des Chasseurs for financial support. Authors are especially thankful to Pascal Lapébie (FNC) for his enthusiasm and scientific input.

## Code availability

The core program is a stand alone written in C. It includes the *lapack* library. Plots were generated using python scripts. All source codes are publicly available at: https://github.com/abdelom/blockbuster_plot

## Extended data

**Table S1:**
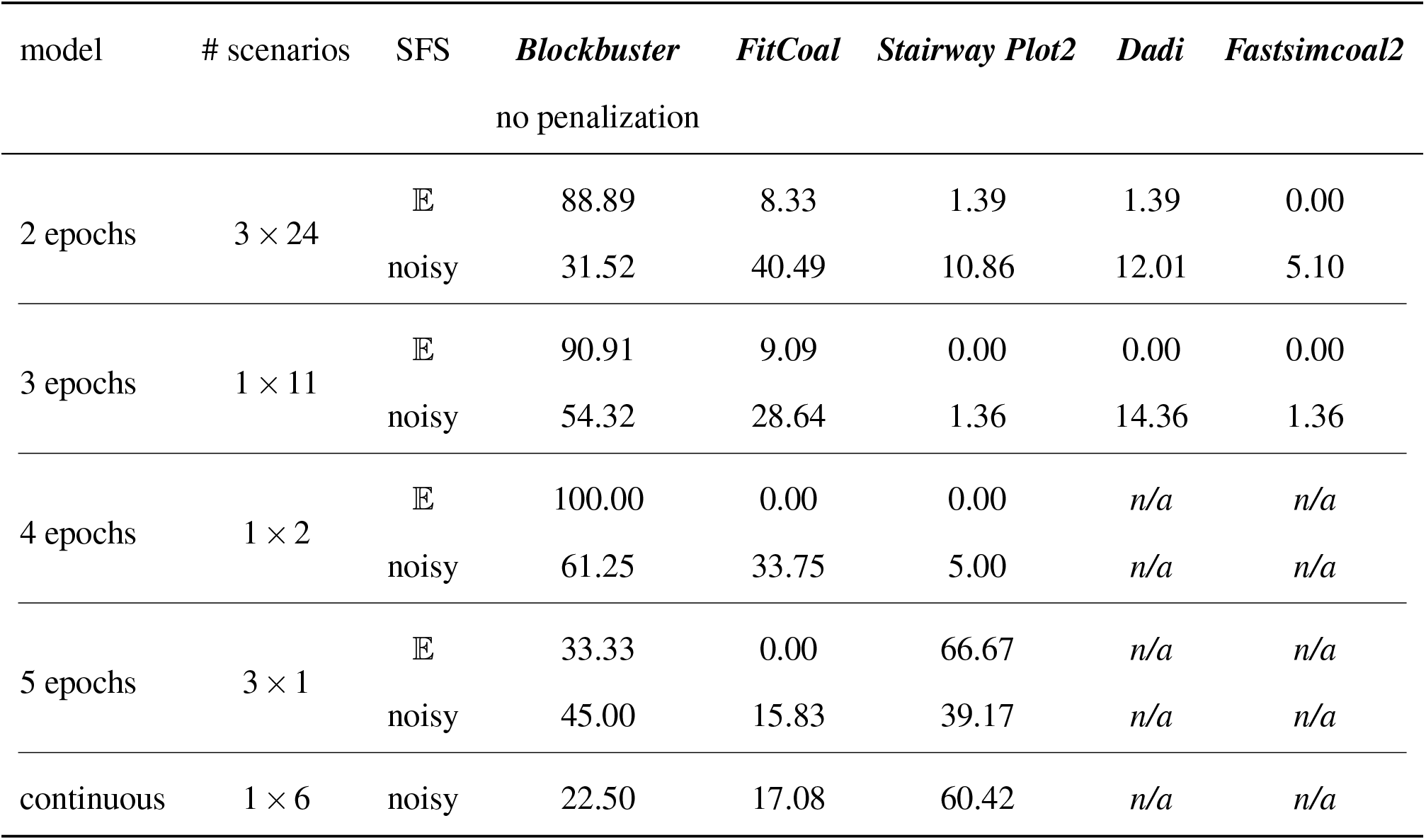
Percentage of inferences for which each program ranks first according to the model category. In each case, results for the expected SFS are shown on top, and results for noisy replicates are shown on the bottom. For partly continuous models only noisy replicates are represented. Here *Blockbuster* does not use likelihood penalization.

**Table S2:** Ranking of programs for the 72 3-parameter models. The top table shows the ranking based on RMSE between inferred and simulated demographic curve for the 72 expected SFS. The bottom table shows the ranking based on RMSE for the 72×40 noisy SFS.

| **Rank** | ***Blockbuster*** | ***FitCoal*** | ***Stairway Plot2*** | ***Dadi*** | ***Fastsimcoal2*** |
| --- | --- | --- | --- | --- | --- |
| 1 | 56.94 | 38.89 | 1.39 | 2.78 | 0.00 |
| 2 | 33.33 | 55.56 | 5.56 | 5.56 | 0.00 |
| 3 | 5.56 | 4.17 | 44.44 | 29.17 | 16.67 |
| 4 | 4.17 | 1.39 | 34.72 | 37.50 | 22.22 |
| 5 | 0.00 | 0.00 | 13.89 | 25.00 | 61.11 |
| 1 | 32.22 | 41.15 | 10.14 | 11.70 | 4.79 |
| 2 | 40.42 | 39.58 | 9.41 | 6.74 | 3.85 |
| 3 | 22.36 | 14.48 | 29.38 | 21.98 | 11.81 |
| 4 | 4.48 | 3.68 | 39.17 | 29.48 | 23.19 |
| 5 | 0.52 | 1.11 | 11.91 | 30.10 | 56.35 |

**Table S3:** Ranking of programs for the 5-parameter models. The top table shows the ranking based on RMSE between inferred and simulated demographic curve for the expected SFS. The bottom table shows the ranking based on RMSE for the noisy SFS (35 replicates).

| **Rank** | ***Blockbuster*** | ***FitCoal*** | ***Stairway Plot2*** | ***Dadi*** | ***Fastsimcoal2*** |
| --- | --- | --- | --- | --- | --- |
| 1 | 100.00 | 0.00 | 0.00 | 0.00 | 0.00 |
| 2 | 0.00 | 72.73 | 9.09 | 9.09 | 9.09 |
| 3 | 0.00 | 27.27 | 18.18 | 54.55 | 0.00 |
| 4 | 0.00 | 0.00 | 36.36 | 27.27 | 36.36 |
| 5 | 0.00 | 0.00 | 36.36 | 9.09 | 54.55 |
| 1 | 62.73 | 25.00 | 1.36 | 9.77 | 1.14 |
| 2 | 28.86 | 42.27 | 7.50 | 13.64 | 7.73 |
| 3 | 7.50 | 24.09 | 21.82 | 42.05 | 4.55 |
| 4 | 0.91 | 6.59 | 39.55 | 26.82 | 26.14 |
| 5 | 0.00 | 2.05 | 29.77 | 7.73 | 60.45 |

**Table S4:**
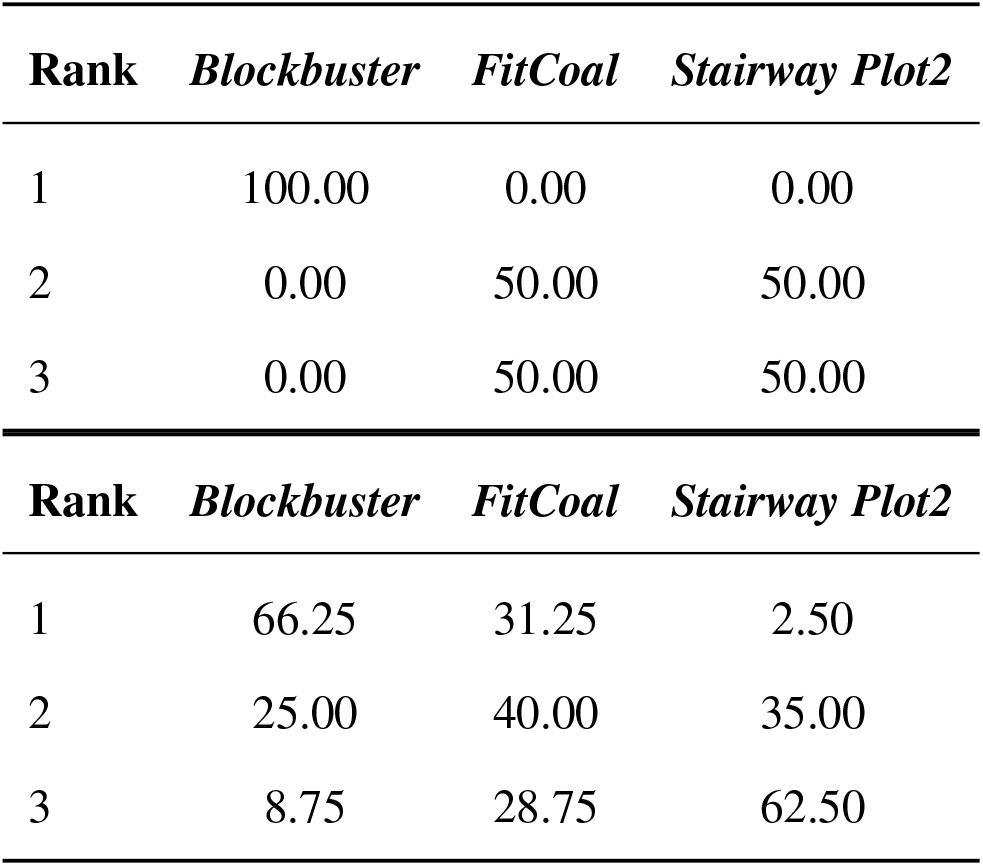
Ranking of programs for the two 4-epoch models. The top table shows the ranking based on RMSE between inferred and simulated demographic curve for the 2 expected SFS. The bottom table shows the ranking based on RMSE for the 2×40 noisy SFS.

| **Rank** | ***Blockbuster*** | ***FitCoal*** | ***Stairway Plot2*** |
| --- | --- | --- | --- |
| 1 | 100.00 | 0.00 | 0.00 |
| 2 | 0.00 | 50.00 | 50.00 |
| 3 | 0.00 | 50.00 | 50.00 |
| 1 | 66.25 | 31.25 | 2.50 |
| 2 | 25.00 | 40.00 | 35.00 |
| 3 | 8.75 | 28.75 | 62.50 |

**Table S5:** Ranking of programs for the two 4-epoch models. The top table shows the ranking based on RMSE between inferred and simulated demographic curve for the 2 expected SFS. The bottom table shows the ranking based on RMSE for the 2×40 noisy SFS.

| **Rank** | ***Blockbuster*** | ***FitCoal*** | ***Stairway Plot2*** |
| --- | --- | --- | --- |
| 1 | 66.67 | 0.00 | 33.33 |
| 2 | 0.00 | 66.67 | 33.33 |
| 3 | 33.33 | 33.33 | 33.33 |

| **Rank** | ***Blockbuster*** | ***FitCoal*** | ***Stairway Plot2*** |
| --- | --- | --- | --- |
| 1 | 82.50 | 4.17 | 13.33 |
| 2 | 14.17 | 20.00 | 65.83 |
| 3 | 3.33 | 75.83 | 20.83 |

**Table S6:** Ranking of programs for the seven continuous linear/exponential size variations. The top table shows the ranking based on RMSE between inferred and simulated demographic curve for the 8x40 noisy SFS (no analytical formula is available under such scenarios for the expected SFS).

| **Rank** | ***Blockbuster*** | ***FitCoal*** | ***Stairway Plot2*** |
| --- | --- | --- | --- |
| 1 | 22.50 | 18.75 | 58.75 |
| 2 | 38.75 | 45.42 | 15.83 |
| 3 | 38.75 | 35.83 | 25.42 |

**Figure S1:**
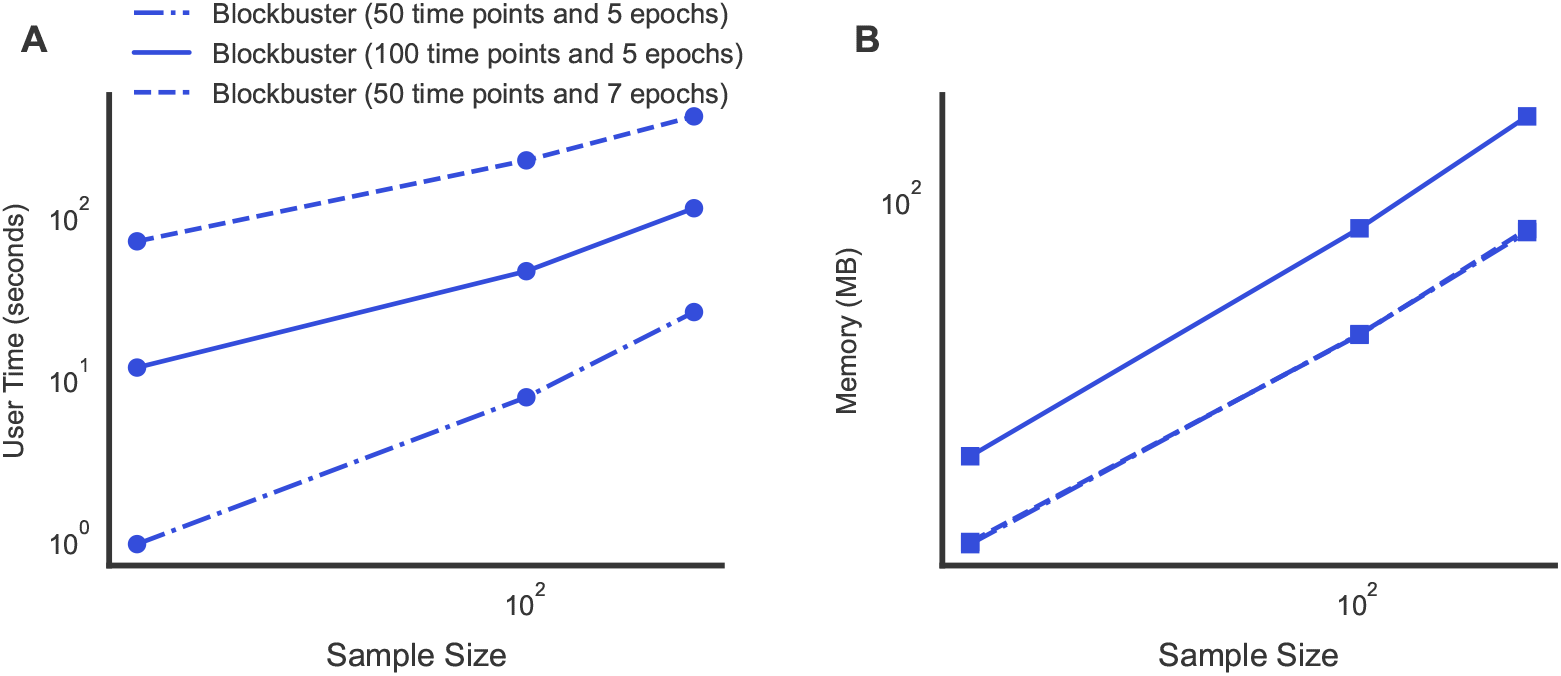
Computational efficiency and memory requirements for 5-epoch demographic inference models. Panels **A–B** assess the computational performance of three *Blockbuster* configurations: up to 5 epochs with 50 (dash-dot) and 100 (solid) time points in the time grid, and up to 7 epochs with 50 time points (dashed).

**Figure S2:**
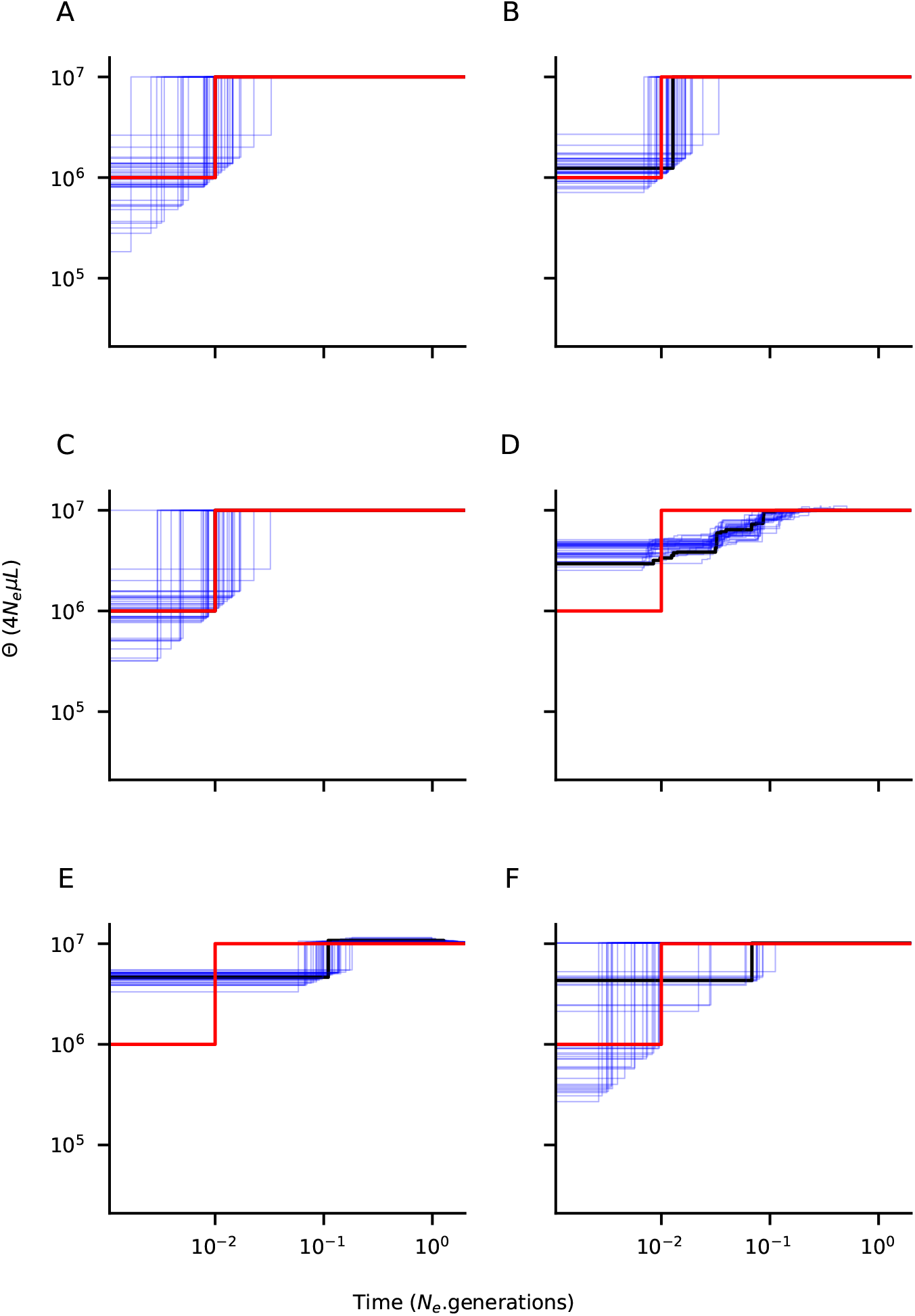
Demographic inference for a simulated two-epoch scenario. The simulated demographic history is shown in red, the inference from the exact expected SFS in black, and the results from 40 noisy SFS replicates in blue. The sample size was set to 20 haplotypes. **(A)** Inference using *Blockbuster* without likelihood penalization, **(B)** Inference using *Blockbuster* with likelihood penalization, **(C)** Inference using *FitCoal*, **(D)** Inference using *Stairway Plot2*, **(E)** Inference using *Dadi*, **(F)** Inference using *Fastsimcoal2*.

**Figure S3:**
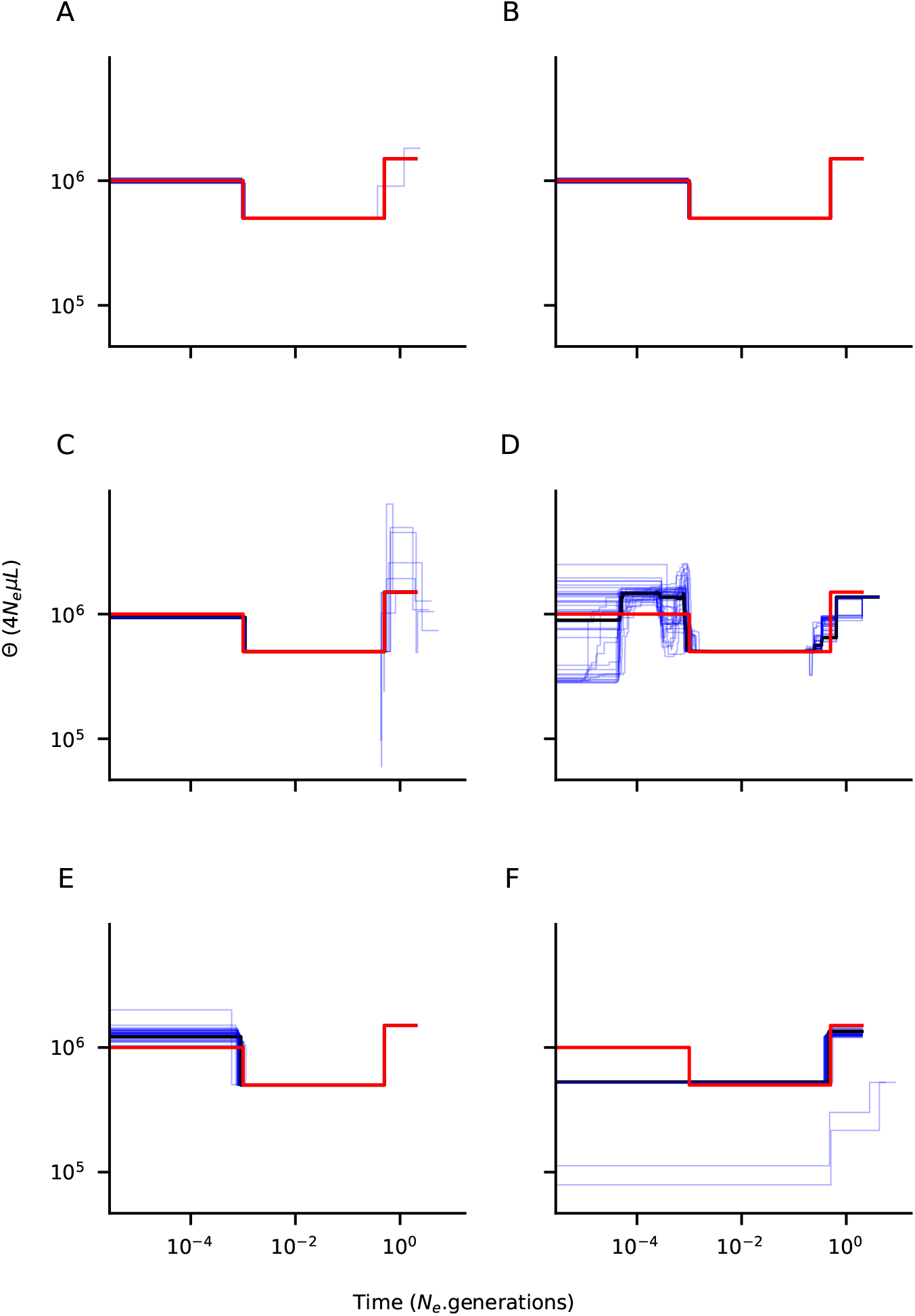
Demographic inference for a simulated three-epoch scenario. The simulated demographic history is shown in red, the inference from the exact expected SFS in black, and the results from 40 noisy SFS replicates in blue. The sample size was set to 500 haplotypes. **(A)** Inference using *Blockbuster* without likelihood penalization, **(B)** Inference using *Blockbuster* with likelihood penalization, **(C)** Inference using *FitCoal*, **(D)** Inference using *Stairway Plot2*, **(E)** Inference using *Dadi*, **(F)** Inference using *Fastsimcoal2*.

**Figure S4:**
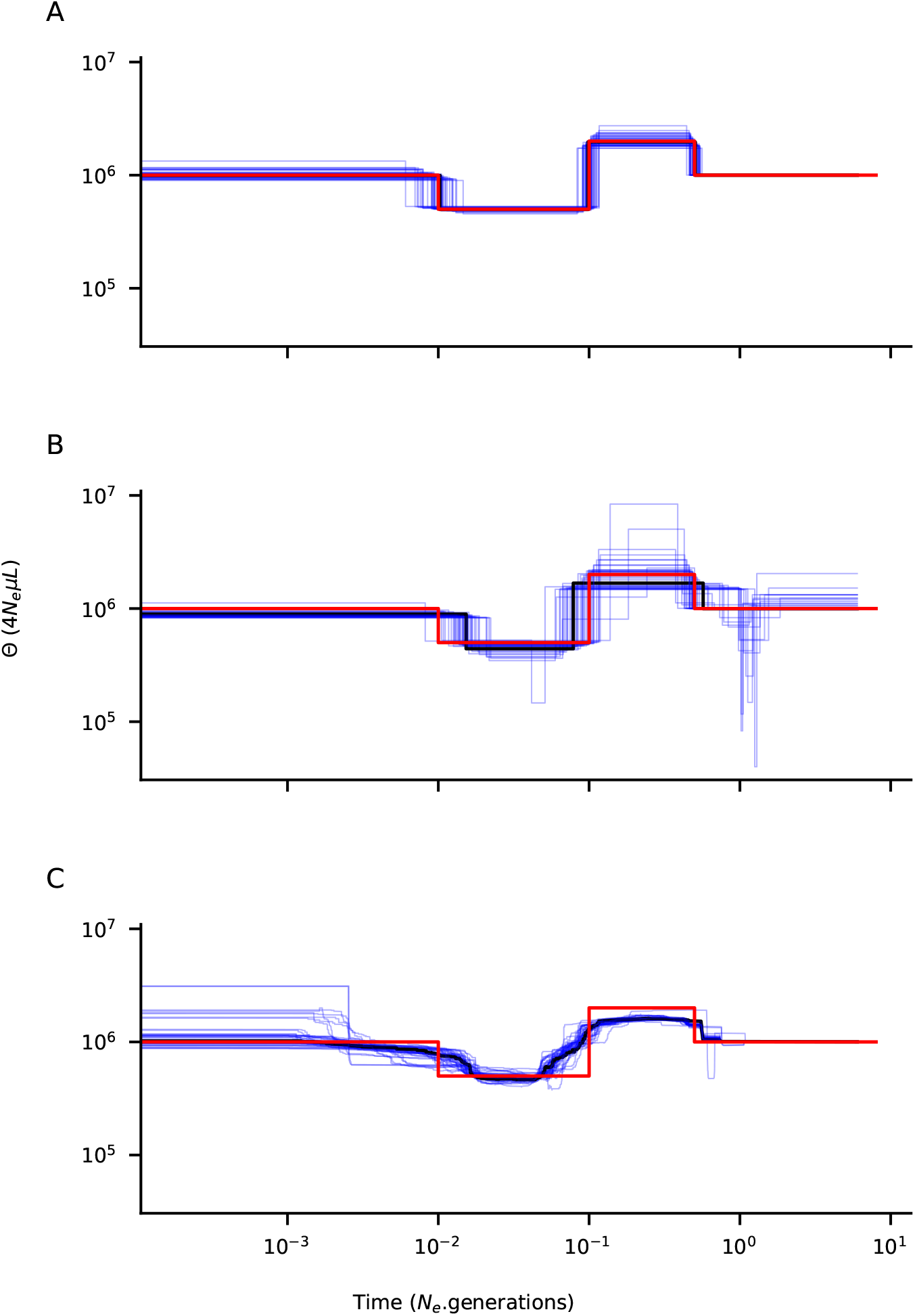
Demographic inference for a simulated four-epoch scenario. The simulated demographic history is shown in red, and the results from 50 noisy SFS replicates are shown in blue. The sample size was set to 50 haplotypes. **(A)** Inference using *Blockbuster* with likelihood penalization, **(B)** Inference using *FitCoal*, **(C)** Inference using *Stairway Plot2*.

**Figure S5:**
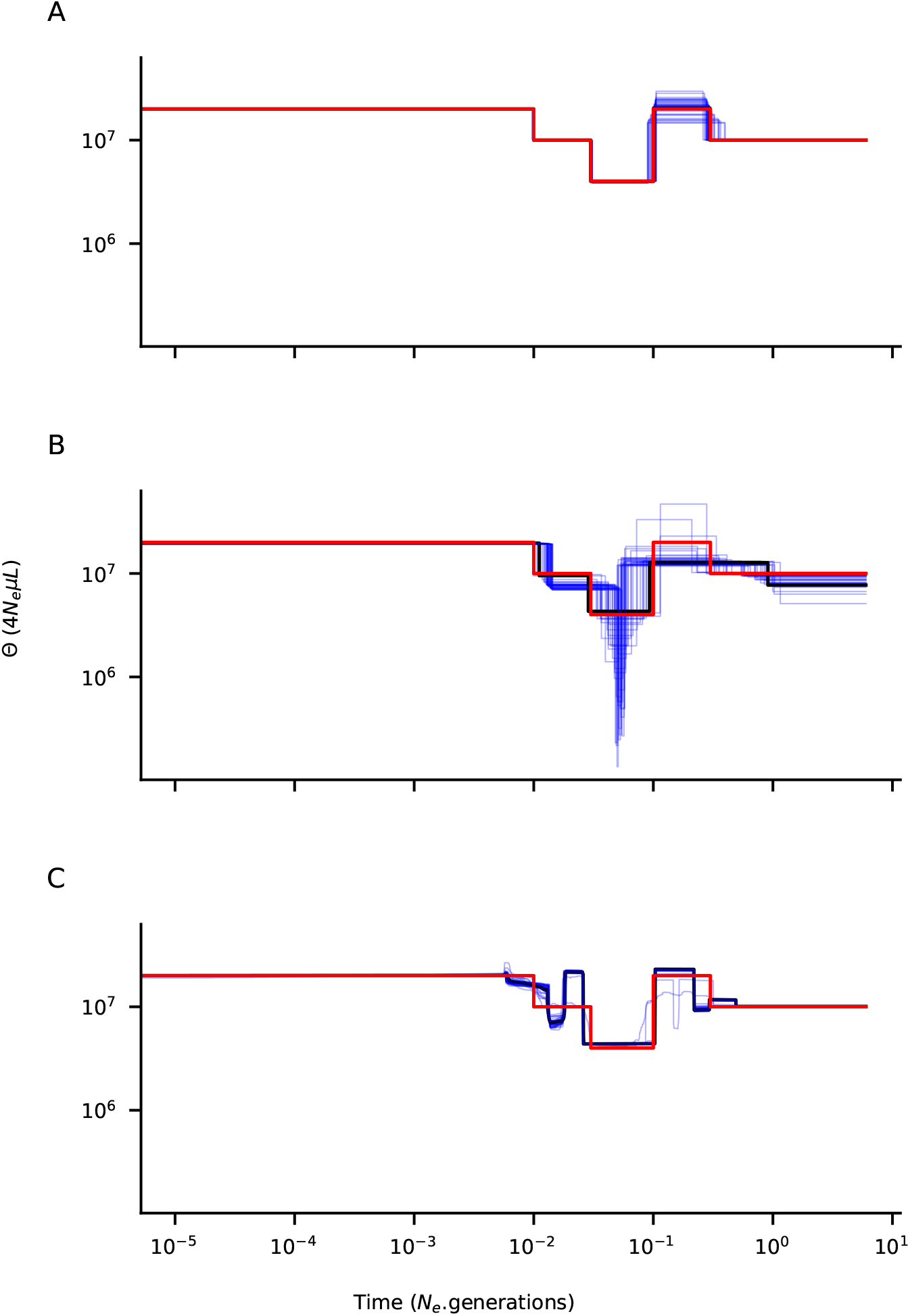
Demographic inference for a simulated five-epoch scenario. The simulated demographic history is shown in red, and the results from 40 noisy SFS replicates are shown in blue. The sample size was set to 100 haplotypes. **(A)** Inference using *Blockbuster* with likelihood penalization, **(B)** Inference using *FitCoal*, **(C)** Inference using *Stairway Plot2*.

**Figure S6:**
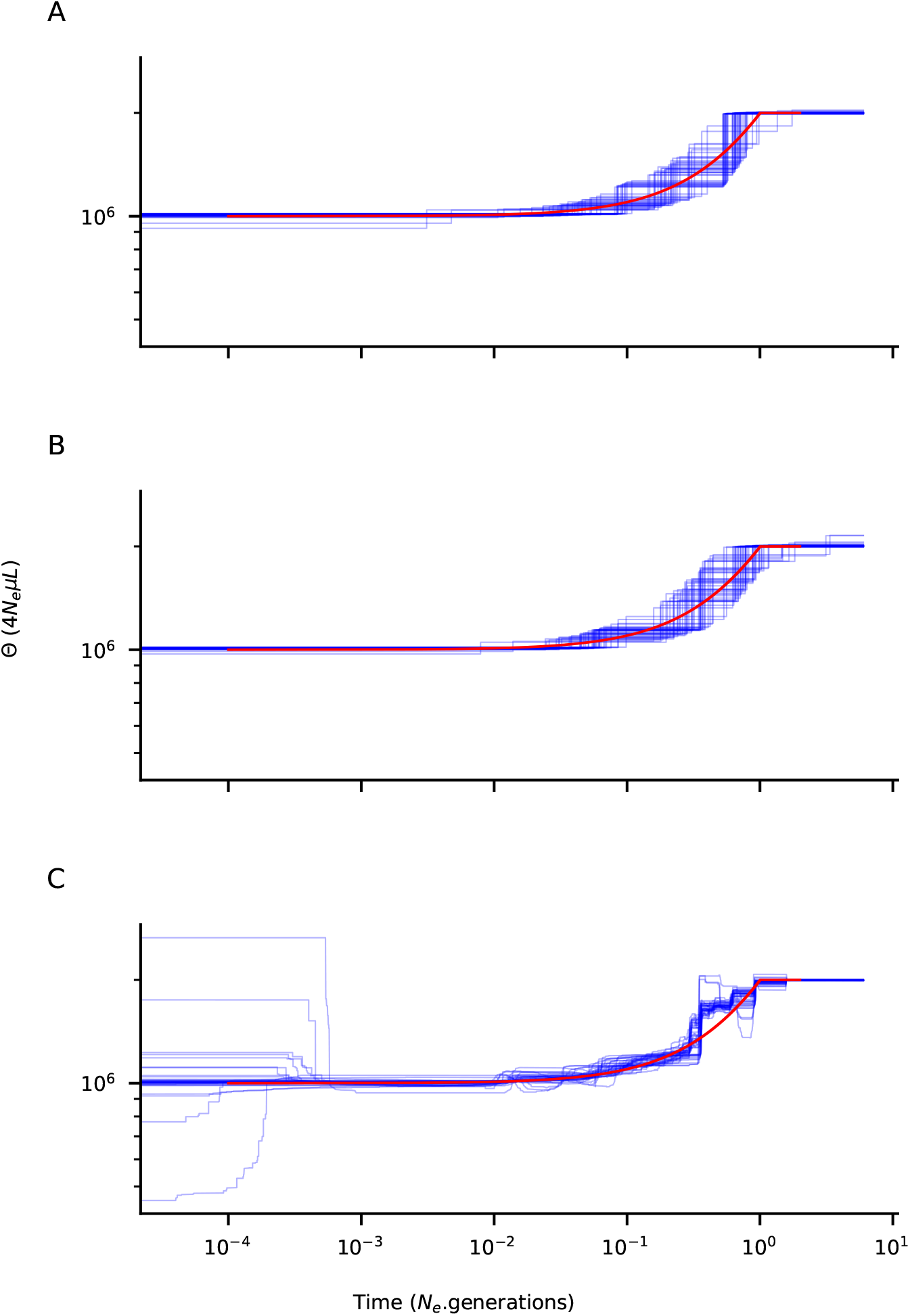
Demographic inference for a simulated scenario with an ancestral plateau, followed by a smooth decline ending with a plateau. The simulated demographic history is shown in red, and the results from 40 noisy SFS replicates are shown in blue. The sample size was set to 100 haplotypes. **(A)** Inference using *Blockbuster* with likelihood penalization, **(B)** Inference using *FitCoal*, **(C)** Inference using *Stairway Plot2*.

**Figure S7:**
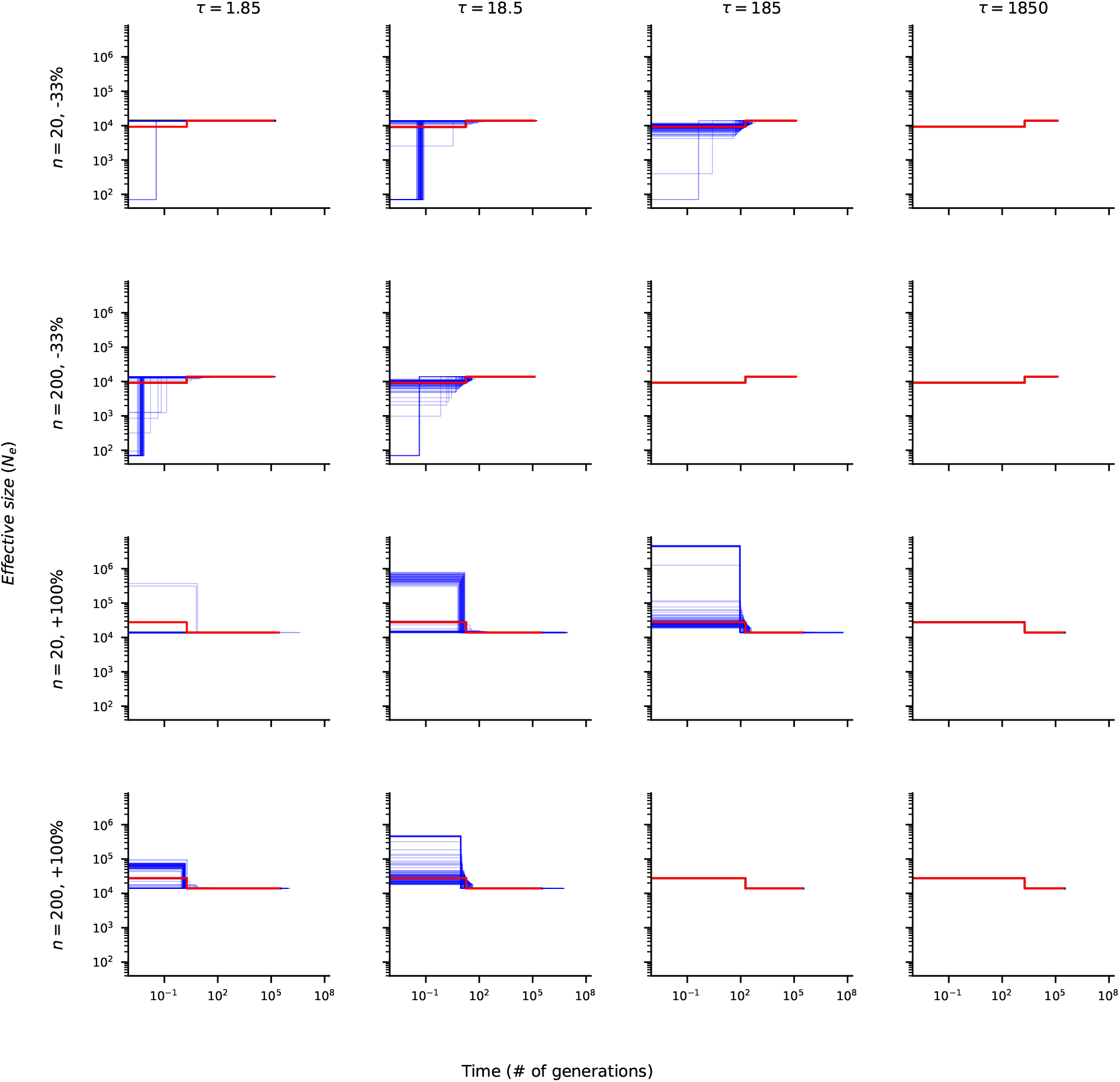
Demographic inference using *Blockbuster without likelihood penalization* for different sample sizes and demographic scenarios, corresponding to either a 33% population decline or a 100% population increase. Subplots show simulated and inferred scenarios for different times of change *τ* ∈ {1.85, 18.5, 185, 1850} and two haploid sample sizes (*n* = 20 and *n* = 200). The simulated demographic history is shown in red, the inferred scenario based on the expected SFS is shown in black, and 100 simulated replicates with sampling noise are shown in blue.

**Figure S8:**
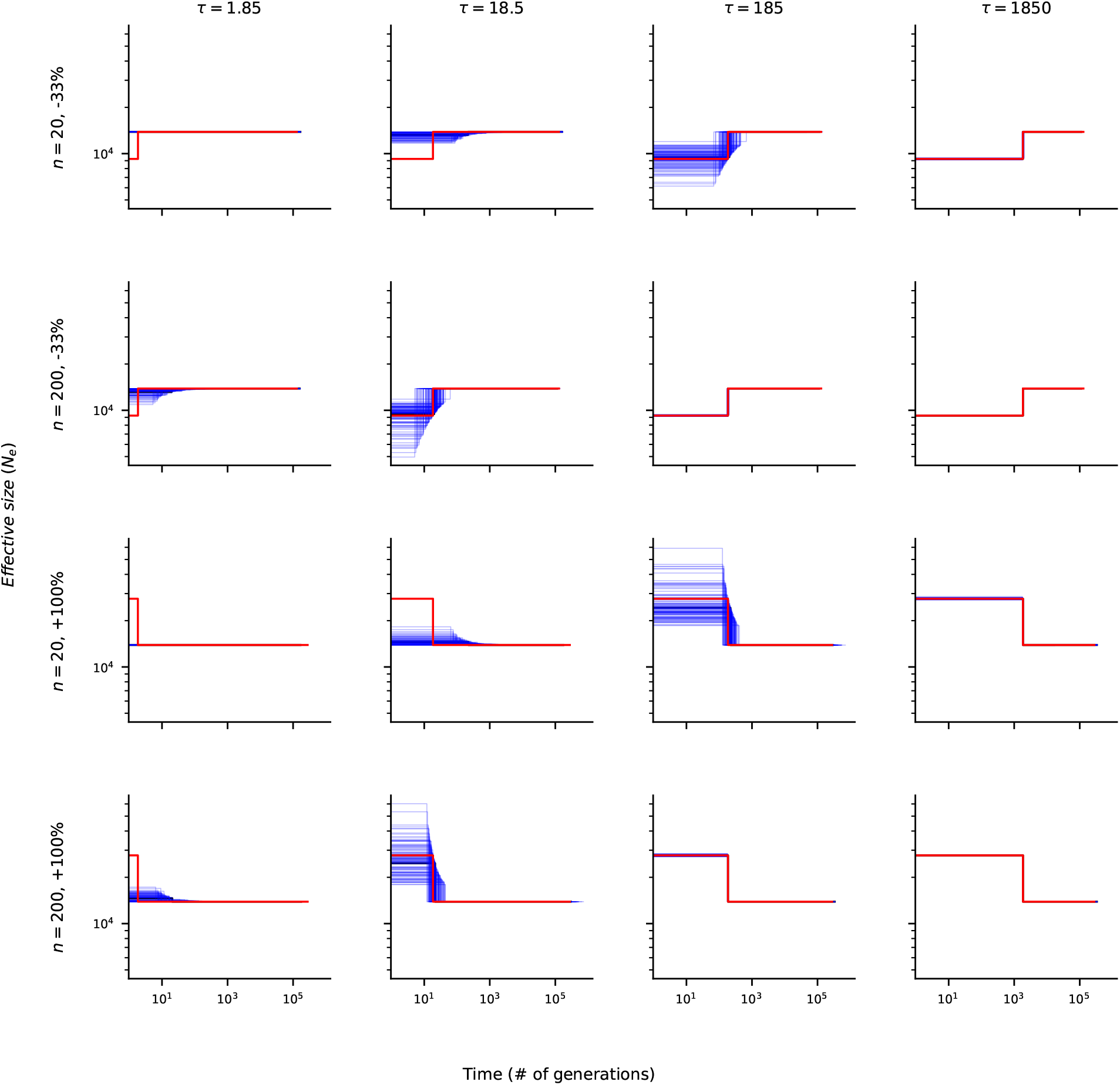
Demographic inference using *Blockbuster with likelihood penalization* for different sample sizes and demographic scenarios, corresponding to either a 33% population decline or a 100% population increase. Subplots show simulated and inferred scenarios for different times of change *τ* ∈ {1.85, 18.5, 185, 1850} and two haploid sample sizes (*n* = 20 and *n* = 200). The simulated demographic history is shown in red, the inferred scenario based on the expected SFS is shown in black, and 100 simulated replicates with sampling noise are shown in blue.

**Figure S9:**
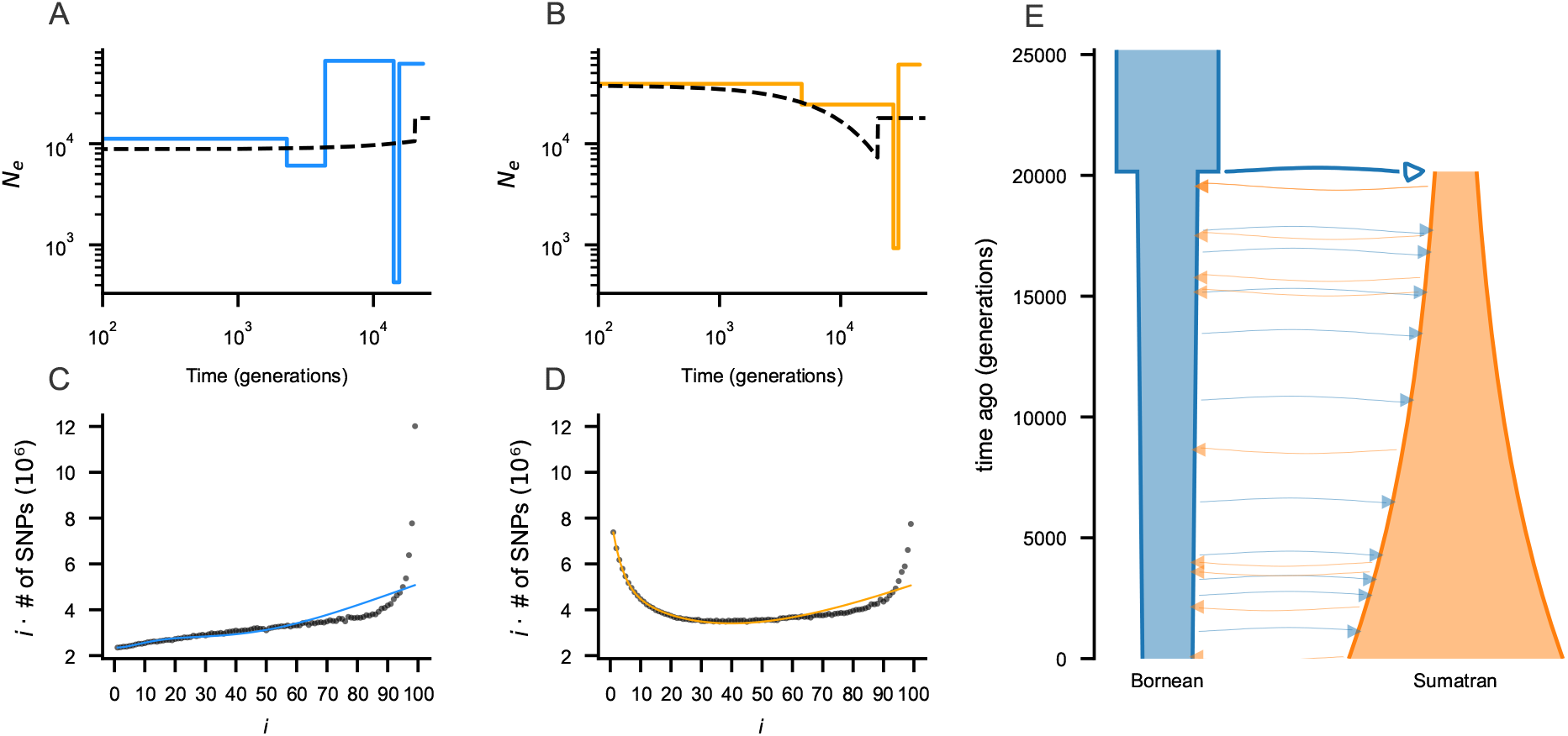
Demographic inference from simulated Site Frequency Spectra (SFS) generated with the popsim library for orangutans. The complete unfolded SFS were used. Panels **A–B** show the demographic histories inferred by *Blockbuster* (colored) from a single simulated SFS of the Bornean (**A**) and Sumatran (**B**) populations; the black curves indicate the true simulated scenarios. Panels **C–D** display the corresponding observed complete SFS (black dots) and the fitted SFS obtained with *Blockbuster* (colored line). Panel **E** shows the joint scenario including both populations with migration.

**Figure S10:**
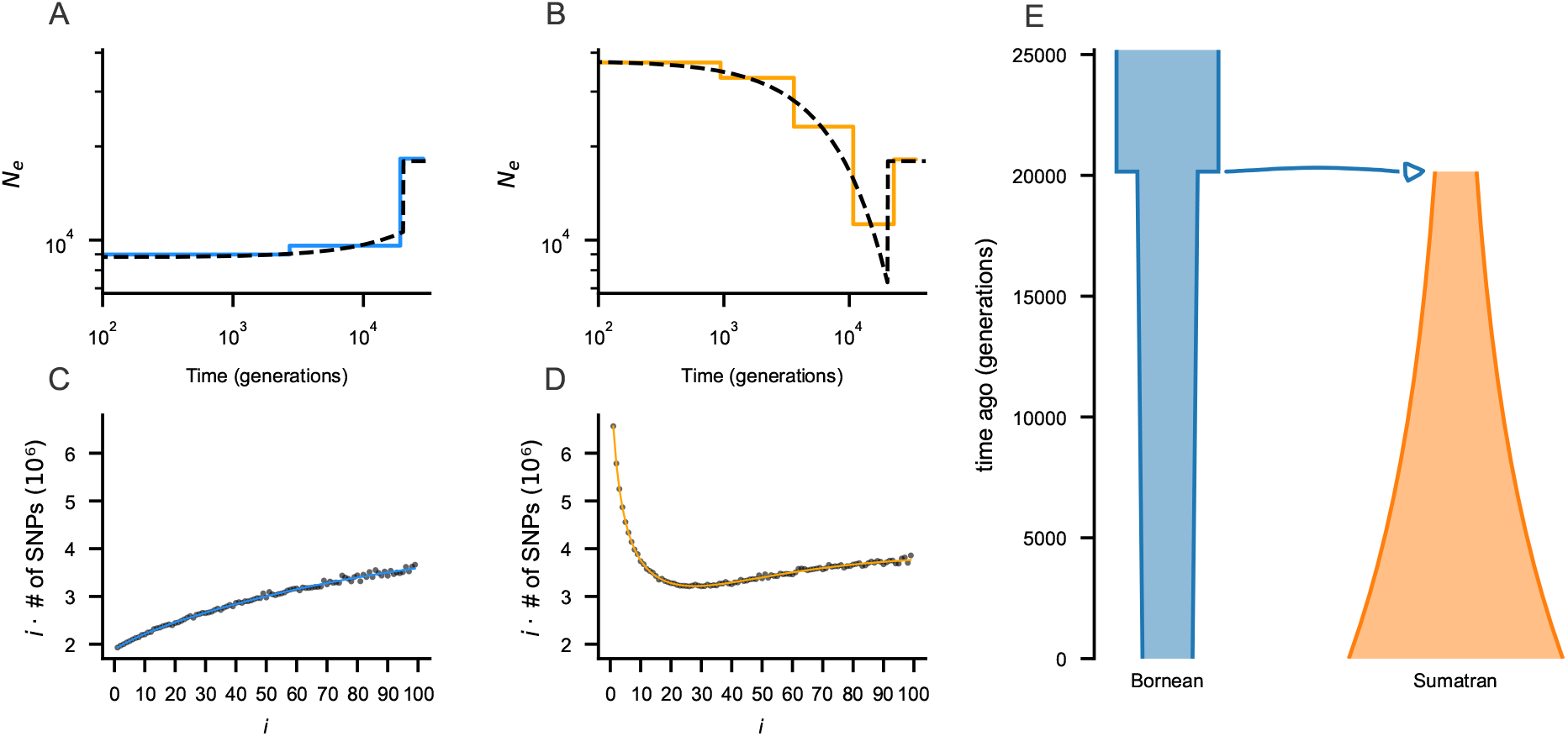
Demographic inference from simulated Site Frequency Spectra (SFS) generated with the pop-sim library for orangutans. The complete unfolded SFS were used. Panels **A–B** show the demographic histories inferred by *Blockbuster* (colored) from a single simulated SFS of the Bornean (**A**) and Sumatran (**B**) populations; the black curves indicate the true simulated scenarios. Panels **C–D** display the corresponding observed complete SFS (black dots) and the fitted SFS obtained with *Blockbuster*, with penalization (colored line). Panel **E** shows the joint scenario including both populations, with migration rate set to zero immediately after divergence and maintained until the present.

**Figure S11:**
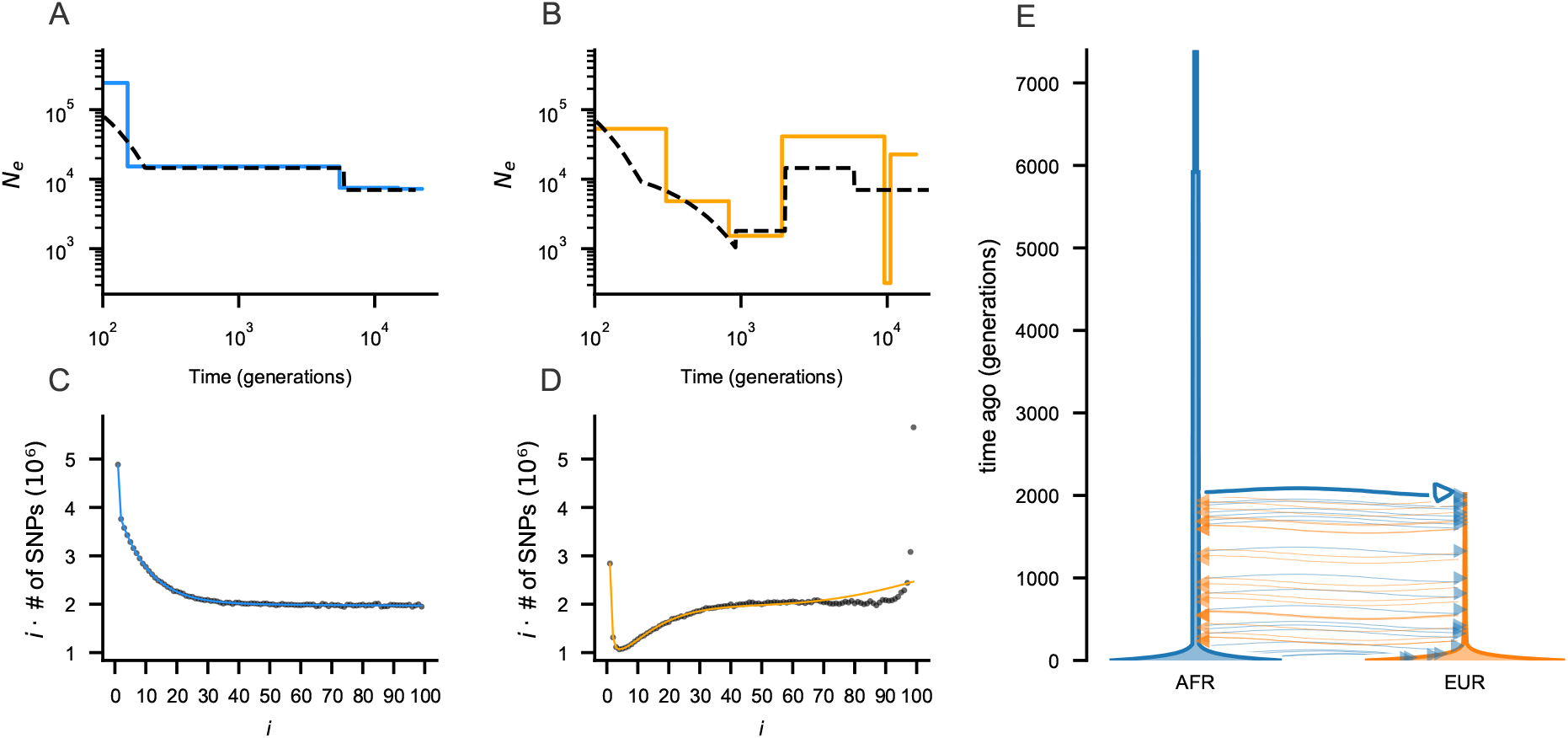
Demographic inference from simulated Site Frequency Spectra (SFS) generated with the pop-sim library for *Homo sapiens*. The complete unfolded SFS were used. Panels **A–B** show the demographic histories inferred by *Blockbuster* (colored) from a single simulated SFS of the African (**A**) and European (**B**) populations; the black curves indicate the true simulated scenarios. Panels **C–D** display the corresponding observed complete SFS (black dots) and the fitted SFS obtained with *Blockbuster*, with penalization (colored line). Panel **E** shows the joint scenario including both populations with migration.

**Figure S12:**
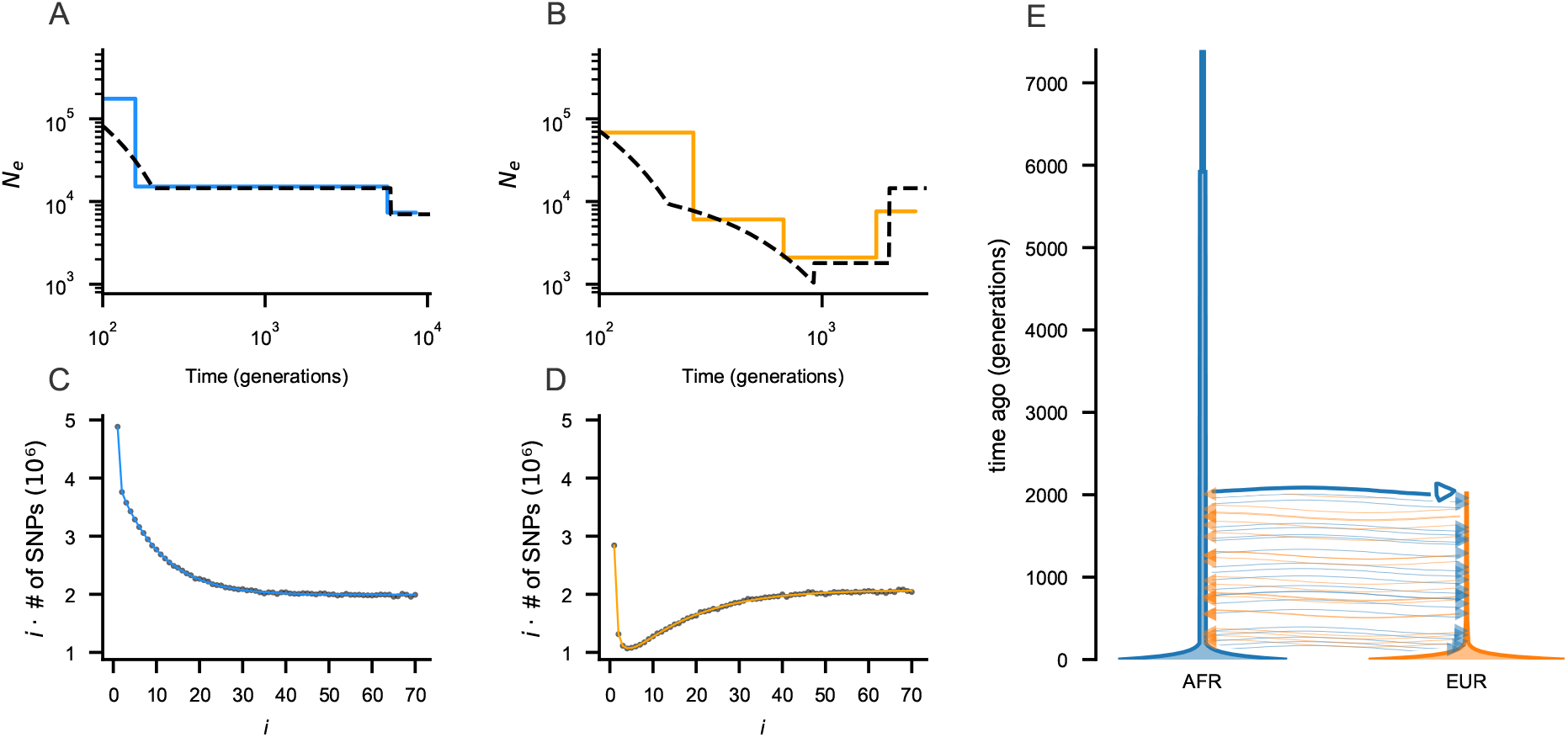
Demographic inference from simulated Site Frequency Spectra (SFS) generated with the pop-sim library for *Homo sapiens*. The complete unfolded SFS were used. Panels **A–B** show the demographic histories inferred by *Blockbuster* (colored) from a single simulated SFS of the African (**A**) and European (**B**) populations; the black curves indicate the true simulated scenarios. Panels **C–D** display the corresponding observed complete SFS (black dots) and the fitted SFS obtained with *Blockbuster*, with penalization (colored line). Panel **E** shows the joint scenario including both populations with migration.

**Figure S13:**
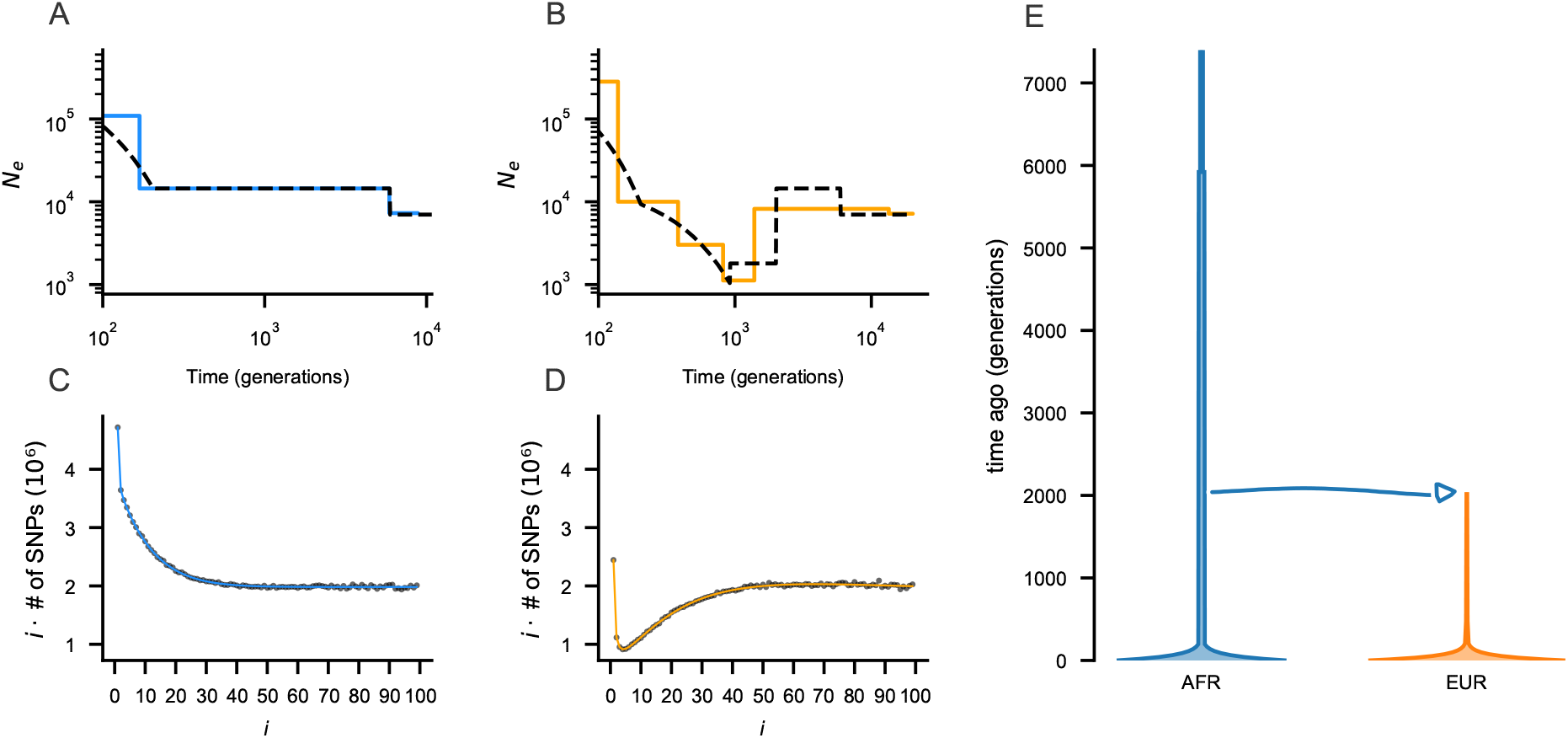
Demographic inference from simulated Site Frequency Spectra (SFS) generated with the pop-sim library for *Homo sapiens*. The complete unfolded SFS were used. Panels **A–B** show the demographic histories inferred by *Blockbuster* (colored) from a single simulated SFS of the African (**A**) and European (**B**) populations; the black curves indicate the true simulated scenarios. Panels **C–D** display the corresponding observed complete SFS (black dots) and the fitted SFS obtained with *Blockbuster* (colored line). Panel **E** shows the joint scenario including both populations, with migration rate set to zero immediately after divergence and maintained until the present.

**Figure S14:**
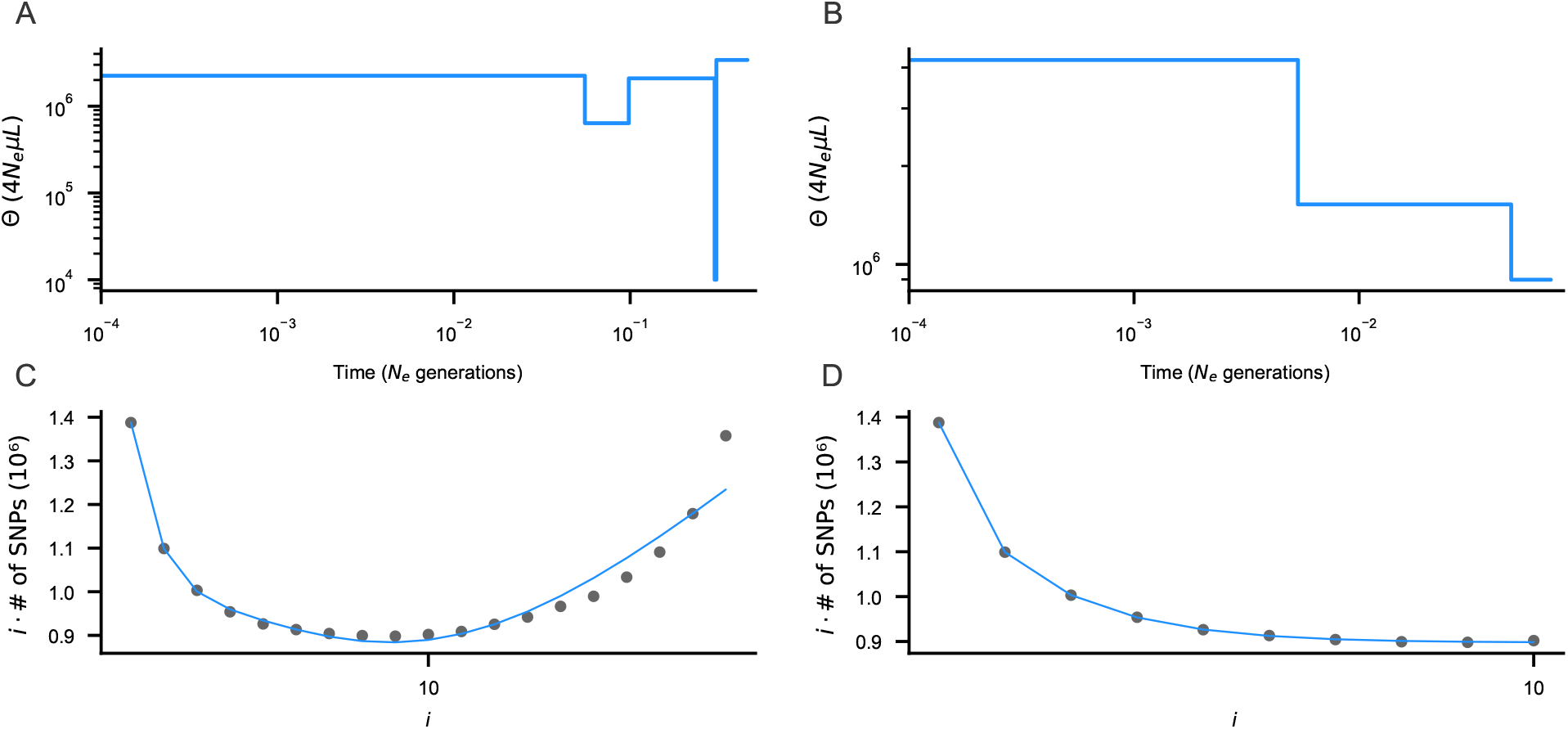
Demographic inference from a single simulated SFS under a Beta-coalescent model with parameter 1.75. Panels **A** and **B** show inferred demographic histories using different input SFS types: (**A**) *Blockbuster* with likelihood penalization, using the complete SFS, (**B**) inference using the truncated unfolded SFS. Panels **C** and **D** show the corresponding transformed observed (black) and fitted (blue) SFS for each inference program, respectively.

## Supplementary Material

### 1 Site Frequency Spectrum under classical coalescent with changing population size

#### 1.1 Basic Maximum likelihood method framework

The expected value of bin *i* (counting variants of frequency *i/n*) in the Site Frequency Spectrum was derived by Fu^1^ as:

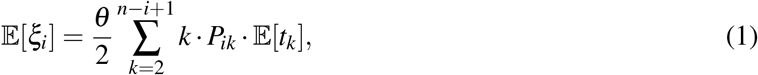

where:

- *θ* = 4*N*_*e*_*µL* is the population mutation rate, with *N*_*e*_ the diploid effective population size, *µ* the mutation rate per site per generation and *L* the genome length (number of sites).
- *k* notes both the stage index in the coalescent tree and count the number of lineages of that stage (*k* ∈ [2, *n*]).
- *P*_*ik*_ is the probability for a given lineage at stage *k* to have *i* descending leaves at the present and is computed as 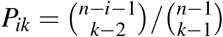.
- *t*_*k*_ is the sojourn time, measured in units of 2*N*_*e*_ generations (for diploids), between two coalescence events and thus delimiting the *k*-th stage in the tree. The law of *t*_*k*_ depends on the demographic scenario, as explained further in the text.

Griffiths & Tavare^2^ developed a framework to simulate expected coalescence times under various population size change scenarios using a Monte Carlo approximation method.

Later, analytical expressions for coalescence times under models with changing population sizes^3,4,5^ were derived, along with methods to make numerical calculations feasible for large sample sizes under piecewise-constant^4^ and exponential growth models^6^.

Assuming that the observed SFS follows a multinomial distribution^7^ with parameters corresponding to the expected SFS bin values expressed as frequencies *f*_*i*_, its composite log-likelihood given the demographic scenario parameters Θ can be written as a sum^8^:

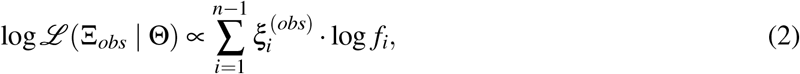

where

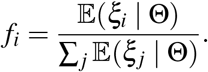

and Ξ_*obs*_ denotes the vector of the observed SFS, with all counts 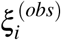 for variants of frequency *i/n* ∈ [1, *n*− 1].

The setup described above enables the use of likelihood maximization methods to fit models such as exponential or linear growth. In the specific case of the piecewise-constant model, where population sizes are assumed constant between instantaneous times of change, an identifiability problem arises between the times and magnitudes of change. In addition, the likelihood surface is rugged for the MC approximation.

Various programs for inferring demographic models have been developed, either based on a backward genealogies^9,10,11,12^ or forward diffusions^13,14^. In the next section, we introduce a popular method based on the backward representation.

#### 1.2 Stairway Plot

Liu & Fu^11^ introduced a program named *Stairway Plot*, which first analyzed unfolded SFS, and was later generalized to the folded spectrum^12^. The authors used a moment estimator for sojourn times, originally applied in the skyline plot for a given tree^15,16^ that coincides the times of change with coalescence events in the genealogical tree(s), such that:

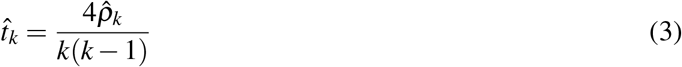

where 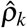 is the ratio of the population size between *t*_*k*_ and *t*_*k*+1_ to *N*_*e*_.

By substituting equation (3) into equation (1) the expected value of bin *i* of the SFS can then be expressed as

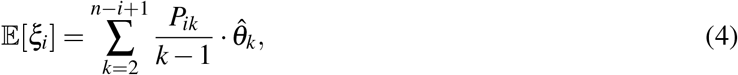

where

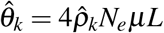

is the so-called population mutation rate at stage *k*.

We note that the *Stairway Plot* approximation assumes that the population sizes of all other stages have no effect on the expected value coalescence time of a given stage *t*_*k*_. A more rigorous (and thus correct) approach is provided in the next section.

Briefly, the algorithm behind *Stairway Plot* is to fix the number and values of times of change and then maximize the SFS likelihood over 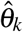, possibly having 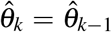. For a given number of changes, the combination of times of change that maximizes the likelihood is retained. The number of changes is progressively increased until the likelihood ratio test fails. The grid of times of change in population size is explored using a heuristic algorithm: a model with *m* changes is constructed from the best solution with *m* − 1 parameters. The additional time is selected from the times that are not yet included in the previous *m*− 1 ones.

The grid is explored using a heuristic algorithm: a model with *m* changes is constructed from the best solution with *m* − 1 parameters, the additional time being selected from those not yet included among the previous *m*− 1.

### 2 An exact linear combination to express the SFS

A more rigorous methodology consists of assuming that the effective population size is constant between pairs of fixed times rather than between pairs of coalescence times. We thus assume that there are times 0 = *H*_0_ < *H*_1_ < · · · *H*_*m*_ < *H*_*m*+1_ = ∞, measured backwards from the present in units of 2*N*_*e*_ generations, such that the effective population size between *H*_*l*_ and *H*_*l*+1_ is constant equal to *N*_*e*_(*l*). Let *ρ*_*l*_ denote the relative effective size between the instantaneous change points *H*_*l*_ and *H*_*l*+1_, defined as

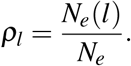

Following previous works^2,3,4,5^, the genealogy under this piecewise-constant model can be viewed as a rescaled version of a constant-size coalescence tree, based on the times

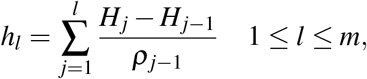

as well as *h*_0_ = 0 and *h*_*m*+1_ = +∞. Using these rescaled times, the expected sojourn times under the piecewise-constant model can be expressed as:

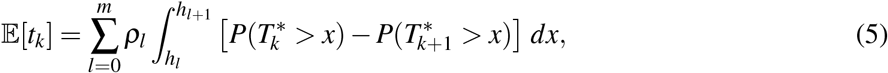

where 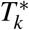 denotes the coalescence time under a constant population size, and *k* ranges from 2 to *n*. For given change points *h*_*l*_ and *h*_*l*+1_, we define the integral term:

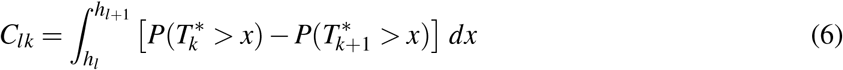

This can be computed as

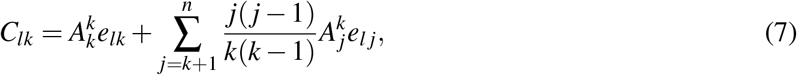

with

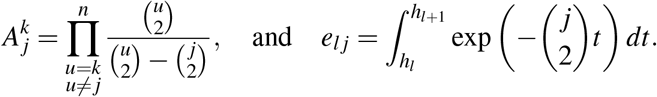

Substituting Equation (5) into Equation (1), we get

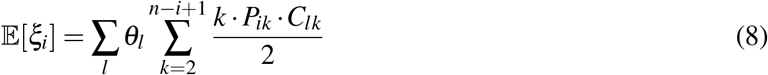

where *θ*_*l*_ = *θρ*_*l*_ = 4*N*_*e*_*ρ*_*l*_ *µL*. It is then possible^6^ to factorize the inner sum so that

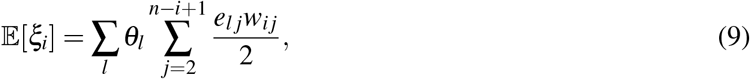

where

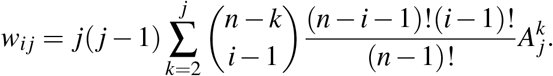

This coefficient can be computed recursively, making the calculation numerically tractable and fast for sample sizes ranging from more than 50 haploid individuals up to 1000, as follows

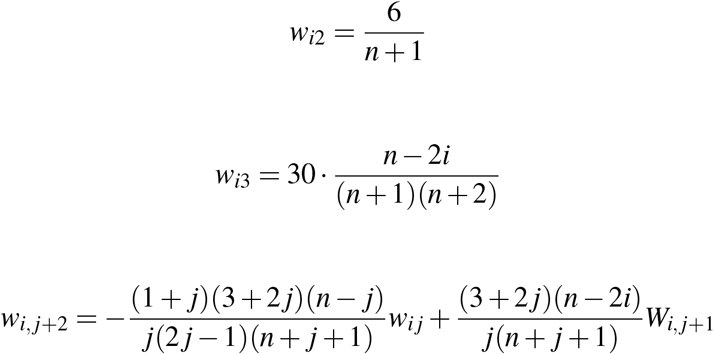

The coefficient associated with *θ*_*l*_ in Equation (9) represents the total branch length subtending *i* individuals in the present, within the considered time interval, under the standard Kingman coalescent with constant population size. Equations (4) and (9) both express each bin of the SFS as a linear combination of the model parameters, allowing for a matrix formulation

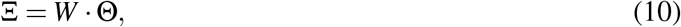

where Θ is the column vector with *i*-th component *θ*_*i*_ and Ξ is the column vector with *i*-th component 𝔼(*ξ*_*i*_ | Θ). The least squares estimator can then be applied as

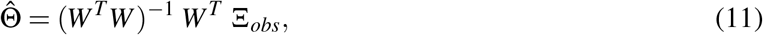

where *W*^*T*^ is the transpose of *W* and Ξ_*obs*_ is the vector with *i*-th component 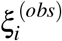. Unlike Equation (4), Equation (8) is not an approximation, but an exact and flexible expression compatible with the Kingman coalescent under varying population size, as developed by^2^. In this framework, time points are no longer constrained to the coalescence nodes of the mean genealogical tree. This approach also closely resembles the simulation-based linear regression method introduced by^17^.

### 3 The *Blockbuster* method

As we just saw, it is possible to analytically compute the population sizes directly from the observed SFS when the times of change are known. We thus propose a likelihood framework in which the only parameters to be optimized are the change points *h*_*l*_ (and then *H*_*l*_).

#### 3.1 Numerical optimisation

##### *Blockbuster* is a three-step algorithm

First, cumulative branch lengths for a given number of leaves in the present are computed over a discrete logarithmic time subdivision of length *H* (number of time points) chosen by the user. This results in a matrix of size (*n* − 1) × *H*, where *n* is the sample size. For a specific number of subtended descendants in the present, these cumulative values are differentiated, in subsequent steps, to obtain the coefficients used in Equation (9), corresponding to the branch lengths within the considered time interval.

Then, an exhaustive search is performed over a coarse sub-sample of the time grid. For each combination of time points explored, the least squares estimator is used to estimate the *θ*_*l*_ values. At the end of the exhaustive search, the best solution is retained.

Finally, the result from the previous step is refined over a second discrete time grid 1000 times larger than the first one and which includes it, using a (very) greedy algorithm: the solution is optimized one time point at a time by selecting the best of its two neighboring values on the grid. The optimization stops when neither neighbor improves the likelihood.

#### 3.2 Model selection

The algorithm described above is applied for a number of instantaneous population size changes ranging from 0 to *Q*, where *Q* is chosen by the user.

The computation time is highly dependent on the parameters chosen by the user. For example, the exhaustive search has factorial complexity and depends on both *Q* and the grid size *H*.

The model that satisfies the likelihood ratio test with two degrees of freedom and a significance threshold of 5% is indicated in the final output.

#### 3.3 Scaling times and sizes

If the user provides neither the mutation rate nor the genome length, then the sizes in the output correspond to temporal population mutation rates, and the times are expressed in units of present-day effective population size generations, i.e., *N*_*e*_ = *N*_*e*_(0). Finally, once the *h*_*l*_ values are obtained, the actual times *H*_*l*_ of change are given by

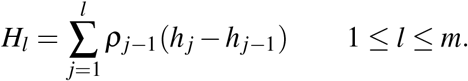

In the case where the user provides both the genome length *L* and the mutation rate *µ, Blockbuster* yields in output the effective size *N*_*e*_(*l*) by dividing the corresponding theta by 4*µL* and the time *H*_*l*_ expressed in number of generations by dividing by the effective size in the present 2*N*_*e*_(0).

Finally, if the generation time is provided, the times are expressed in years.

#### 3.4 Penalization

Similarly to other methods^18,11,12^, we add a penalized mode in *Blockbuster*, in which the likelihood is penalized by the *L*^1^ norm of log-ratios of adjacent *θ* values in a k-epoch model:

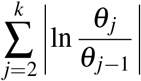

This penalty term has the advantage of being symmetric for both growth and decline phases. It constrains relative changes in effective population size when the likelihood surface is largely flat. As a result, in cases where identifiability issues may arise^19^, it favors moderate changes over strongly discontinuous ones.

### 4 The theoretical limit to the detection of one very recent change

In this section, we seek to understand when an observed SFS generated under a model with a recent demographic change (hypothesis *H*^*′*^) and the SFS without demographic change (hypothesis *H*) can be distinguished statistically. We give analytical formulae for the threshold numbers of generations 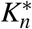 (resp.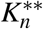) after which the change can be detected (resp. can be estimated). These theoretical lower bounds are independent of the methods designed here or elsewhere. A method that can detect/estimate correctly a change at a date which is as recent as these thresholds can be deemed optimal.

We let *N* be an arbitrary population size serving as unit of measure and consider the following two scenarios:

- Under hypothesis *H*^*′*^, we consider a two-epoch model, where the population size is constant equal to *N*_*pres*_ = *ρ*_*pres*_*N* in recent time and jumps to *N*_*anc*_ = *ρ*_*anc*_*N* at Δ units of 2*N* generations in backward time. The expected value of the *i*-th SFS bin under *H*^*′*^ is denoted by *E*_Δ_(*ξ*_*i*_);
- The SFS obtained under *H* (constant population size equal to *N*_*anc*_ throughout the entire demographic history) has expected value of the *i*-th SFS bin denoted by *E*_∅_(*ξ*_*i*_).

We set as previously *ρ*_*i*_ = *N*_*i*_*/N* and *θ*_*i*_ = *ρ*_*i*_*θ*, where *θ* = 4*NµL, µ* is the mutation probability per generation and per site and *L* is the number of sites. In the next proposition, we compute the expected distance between the two SFS, globally and at each bin separately.

#### Proposition 4.1.

*If the recent demographic change occurred* 2*N*Δ *generations ago, then the L*^1^ *distance between the two SFS can be expressed when n is large, as*

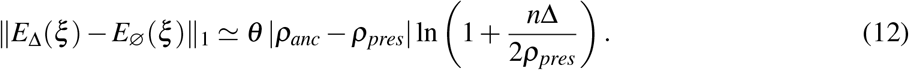

*For each* 1 ≤ *i* ≤ *n*− 1,

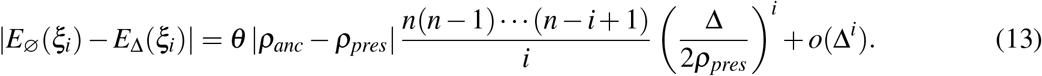

*At the second order in* Δ, *we have*

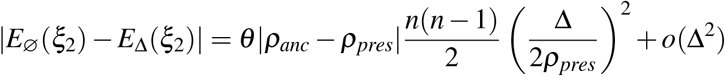

*and*

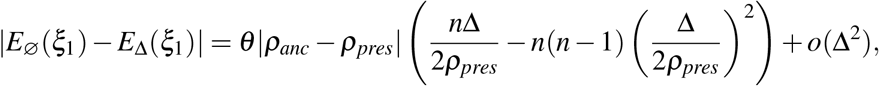

*so that, using* (13),

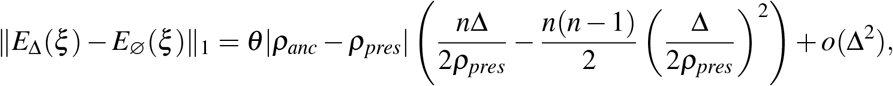

*which is consistent with* (12) *(replacing n*(*n*− 1) *with n*^2^).

#### Remark 4.2.

*Check that* |*E*_∅_(*ξ*_1_) − *E*_Δ_(*ξ*_1_)|*/*∥*E*_Δ_(*ξ*) − *E*_∅_(*ξ*)∥_1_ *converges to 1 as* Δ → 0, *that is, the discrepancy between the two SFS when* Δ *is small is only due to singletons*.

*Proof*. We start with a coupling of the two coalescents *T* and *T* ^*′*^ on a same space. We assume that *N*_*pres*_ < *N*_*anc*_. First, because the topology of the coalescent always has the same law, independent of the demographic scenario, we can assume that *T* and *T* ^*′*^ have the same labelled topology, for example the labels of the first two leaves to coalesce are the same in *T* and in *T* ^*′*^. Then the law of *T* (resp. *T* ^*′*^) is characterized by its coalescence times *τ*_*n*_ < · · · < *τ*_2_ (resp.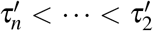). Set *σ*_*i*_ := *τ*_*i*_ − *τ*_*i*+1_ and 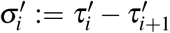, with 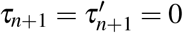.

We first construct the (*τ*_*i*_) by drawing independent random variables (*σ*_*i*_), where *σ*_*i*_ is exponentially distributed with parameter *γ*_*i*_ given by

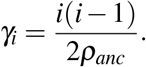

Then we can assume that the (*τ*_*i*_) are given and we now construct jointly the 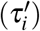 as follows. Set

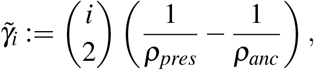

and define the function

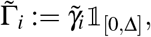

where for any interval *A*, 𝟙_*A*_(*t*) = 1 if *t* ∈ *A* and 0 otherwise. In particular, notice that

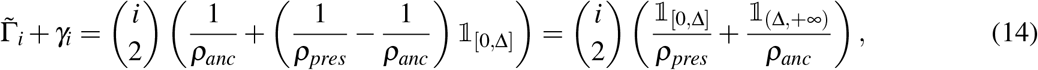

is the inhomogeneous coalescence rate under *H*^*′*^.

We define recursively the 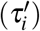 as follows, starting from 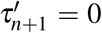. Assume that we have constructed 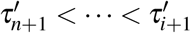. Let 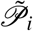 be an inhomogeneous Poisson point process with intensity 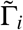, independent of the construction done so far. Then we define

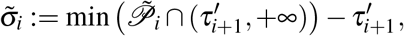

with the convention that min∅ = +∞, and

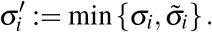

As a consequence, 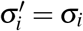 as soon as 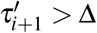 (because the intensity of 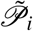 is zero beyond Δ). In particular, *T* = *T* ^*′*^ whenever 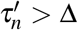. This finishes the recursion because from the definition of *σ*_*i*_, we can write the same representation for *σ*_*i*_ as for 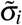

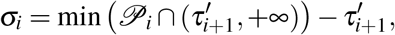

where 𝒫_*i*_ is an independent, homogeneous Poisson point process with intensity *γ*_*i*_, so that

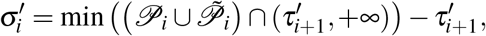

and

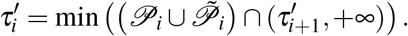

This shows that 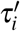 has the desired density, as it is the first point larger than 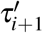 of the Poisson point process 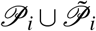, which has inhomogeneous intensity 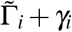 given by Eq (14).

Now let us show that conversely, we can construct the *τ*_*i*_ assuming the 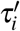 are given. It is well-known that each point *t* in 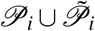 has a probability

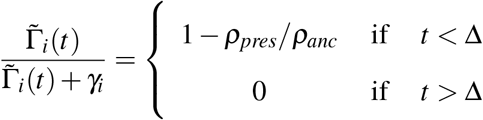

to be in 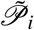 rather than in 𝒫_*i*_, and these events are independent. Then let *ε*_*i*_ be the indicator of the event that the first point in 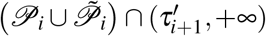 is in 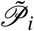. If *ε*_*i*_ = 0, then 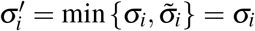. Conditional on 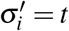 and *ε*_*i*_ = 1 (which can only happen if *t* < Δ), *σ*_*i*_ is an exponential random variable with parameter *γ*_*i*_ conditioned on being larger than 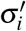, which is equivalent to saying that *s*_*i*_ = *σ*_*i*_ −*t* is exponential with the same parameter. As a consequence we can assume the 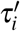 are given and define

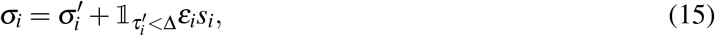

where *s*_*i*_ is exponentially distributed with parameter *γ*_*i*_ and *ε*_*i*_ is an independent Bernoulli random variable with success probability 1 −*ρ*_*pres*_*/ρ*_*anc*_.

Finally, we see that *T* can be obtained from *T* ^*′*^ by simply adding to the time interval of length 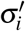 in *T* ^*′*^, for every *i* such that 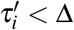, an independent extra length *ε*_*i*_*s*_*i*_. We now study the effect of these differences on the SFS when Δ is small.

Define

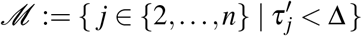

as the set of coalescence events of *T* ^*′*^ that occur before time Δ.

Then it is easily seen that for each *i* = 0, 1, …, *n*− 1,

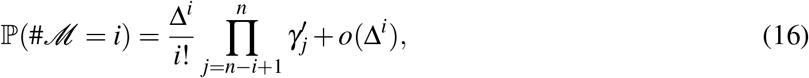

with

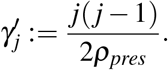

Also, an easy consequence of Equation (15) is that conditional on #ℳ = *i*, the number of *j* such that 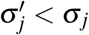 is binomial with parameters *i* and 1 − *ρ*_*pres*_*/ρ*_*anc*_. An even simpler consequence of (15) is that for each *i* ∈ ℳ,

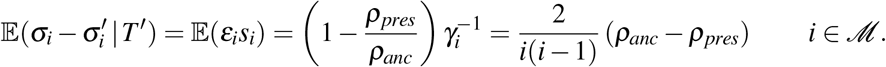

Taking expectation with respect to the law of *T* ^*′*^ yields for any *i* ≥ 2

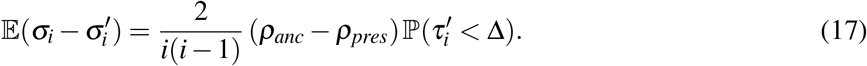

Now we denote by *ℓ*_*k*_ (resp. 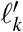) the total length of *T* (resp. *T* ^*′*^) subtending *k* leaves. Recall that for any 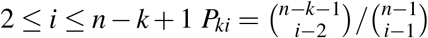 is the probability that a given lineage at stage *i*, i.e., for times in (*τ*_*i*+1_, *τ*_*i*_), has *k* descending leaves. Note that *P*_1*n*_ = 1, *P*_1,*n*−1_ = (*n*− 2)*/*(*n*− 1) and *P*_2,*n*−1_ = 1*/*(*n*− 1). Then

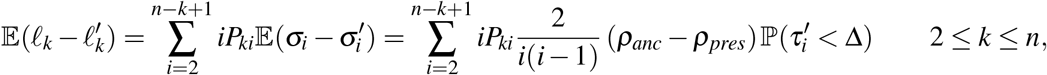

which by changing indices and multiplying by *θ/*2 (*θ* = 4*NµL*, the number of mutations per generation is *µL* and the time unit is 2*N* generations, so the mutation rate per unit time is *θ/*2), becomes

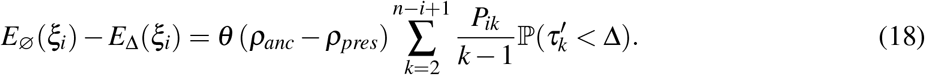

In particular, thanks to (16),

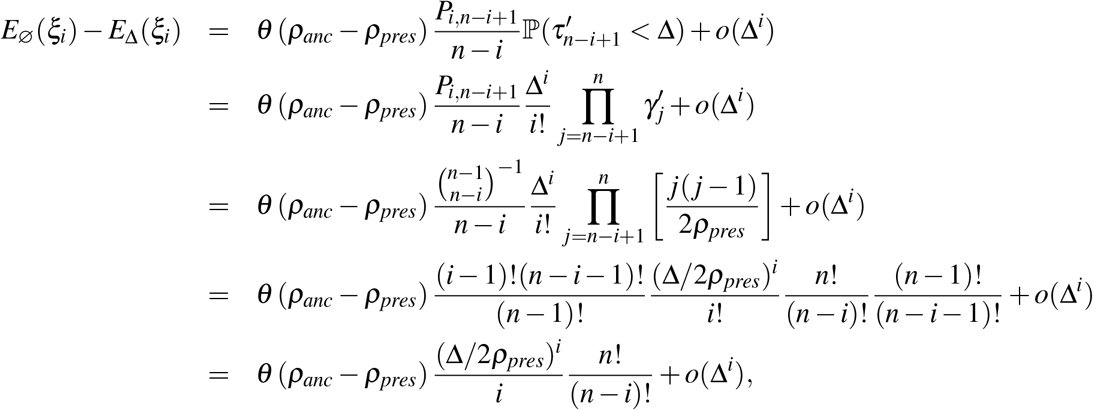

which yields Equation (13) in the proposition. We now prove the next two equations of the proposition. First,

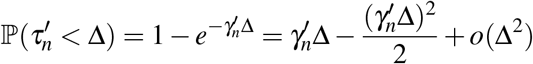

and

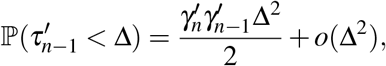

so that, using (18),

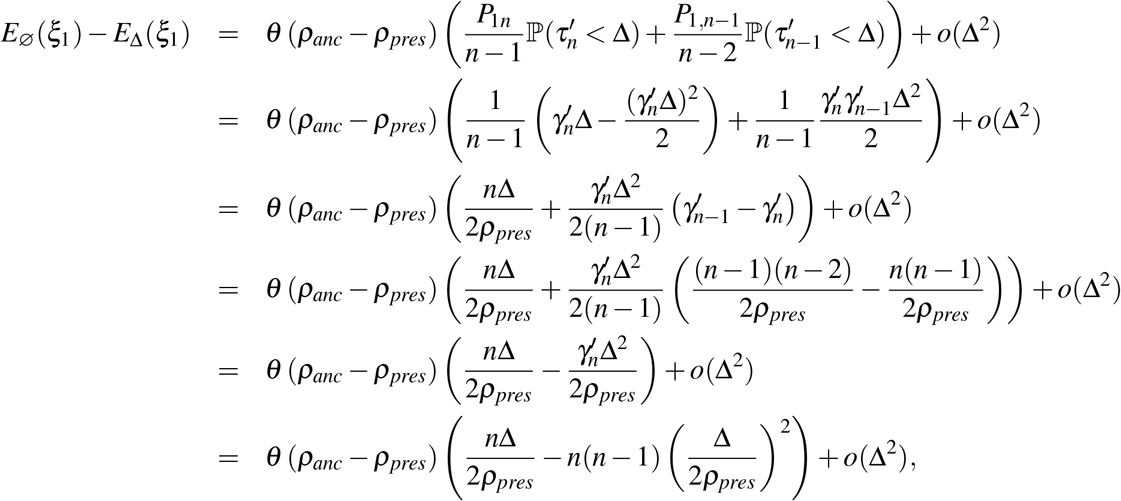

and

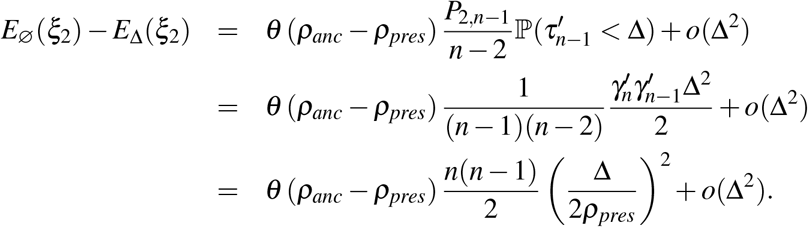

We finally seek to prove the first formula of the proposition. We recall that

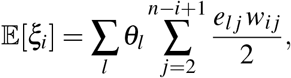

where

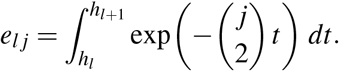

Therefore, we can express the expected SFS as follows:

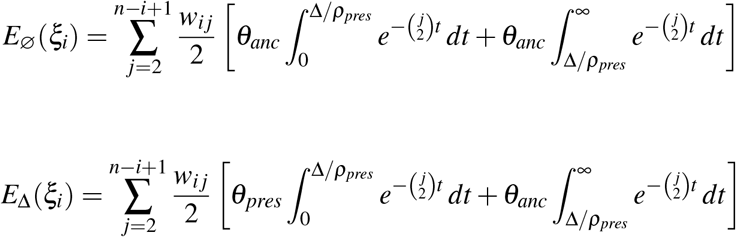

Hence

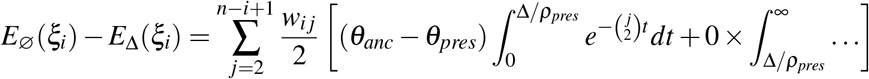

As a consequence, we can compute the *L*^1^ distance between the two SFS as

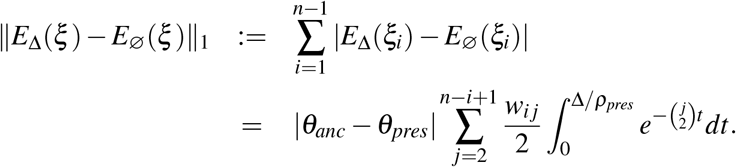

Now this last quantity is the expected sum of all bins of a SFS with piecewise-constant demography, namely with populational mutation rate equal to |*θ*_*anc*_ −*θ*_*pres*_| for times smaller than Δ*/ρ*_*pres*_ and equal to 0 for larger times. The expected sum of bins of a SFS is the total length of the tree times half the populational mutation rate, so that, denoting by *N*_*t*_ the number of lineages at time *t*,

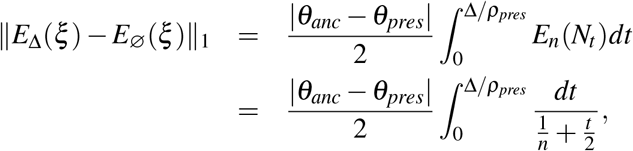

where the last equality is due to the fact that *E*_*n*_(*N*_*t*_) can be approximated when *t* is small (and *n* is large) by the solution to 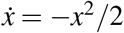 with initial condition *x*(0) = *n*, which is equal to 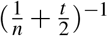. This yields

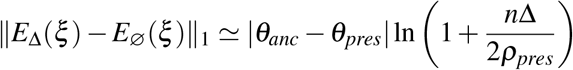

which as Δ → 0, is equivalent to *n*|*θ*_*anc*_ −*θ*_*pres*_| Δ*/*(2*ρ*_*pres*_).

The next two propositions (Propositions 4.4 and 4.5) state that the demographic change is statistically detectable if the number of generations elapsed since it occurred is larger than 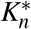 times some positive constant which essentially lies between 1 and 3, where

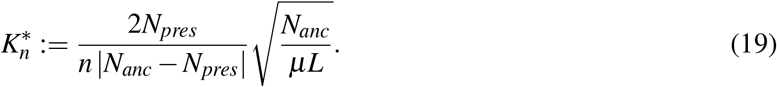

Note that the date 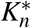 of the most recent detectable demographic change increases like the square root of ancient population size *N*_*anc*_, the inverse of |*N*_*anc*_*/N*_*pres*_ − 1| and the inverse of sample size *n*.

#### Remark 4.3.

*If the data available are whole genome sequences, then µL is the number of genome-wide mutations per generation. Let us assume that µL* = 2 *(a lower bound for most organisms) and that N*_*pres*_ = *N*_*anc*_*/*2 *(population halving), then the threshold number of generations is* 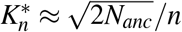.

#### Proposition 4.4

(Type I error). *The probability of rejecting H*^*′*^ *under H, that is, of not attributing a deficit (resp. excess, resp. deviation) of singletons to a false population decline (resp. expansion, resp. change), k generations ago, is larger than w if*

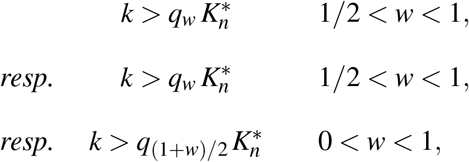

*where q*_*w*_ *is the standard normal w-quantile, such that*

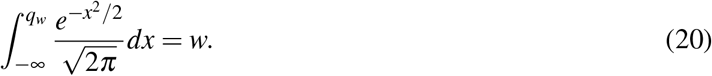

*Proof*. We wish to determine whether the noise inherent to real SFS measurement in bin 1 can be erroneously attributed to a recent demographic change. For simplicity, assume that we can chop the genome into *M* chunks of equal length *ℓ* that can be considered as independent realizations of the standard SFS with populational mutation rate 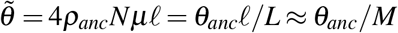. Let us denote by *X*_*j*_ the number of singletons in the *j*-th chunk, so that 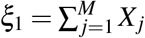. When *L* (and so *M*) is large, under hypothesis *H*, 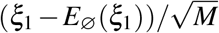 behaves like a centered normal random variable with variance 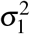 given by^1^

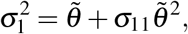

where

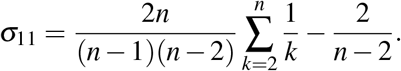

As a consequence, for any *w* > 1*/*2, the probability of a type I error, i.e., of inferring wrongly that a deficit of singletons (compared to the expected standard SFS) is due to a population decline, is smaller than 1 −*w* if

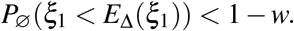

Now the left-hand side is

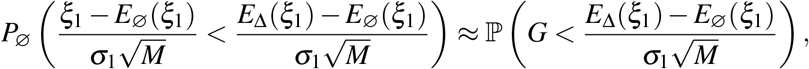

where *G* is a standard normal random variable. As a consequence,

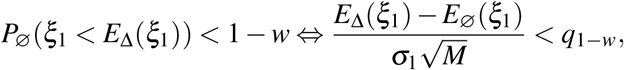

which is equivalent to

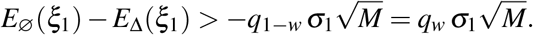

Replacing the left-hand side with its expression yields

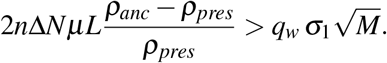

Now we can pick *ℓ* sufficiently small that 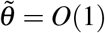 and so 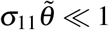, since *σ*_11_ only depends on *n* and goes to 0 as 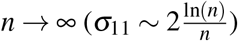. Then we have 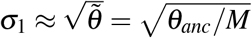 and the condition becomes

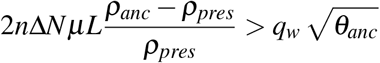

or equivalently

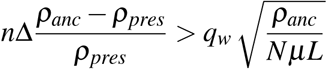

When 2*N*Δ = *k* generations, this is equivalent to

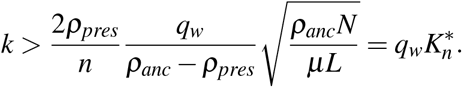

The probability of the type I error consisting in inferring wrongly a deviation in the number of singletons (compared to the expected standard SFS) can be obtained in the same way, replacing differences between expected singleton frequencies with their absolute value, and the one-sided test with a two-sided test.

In the following proposition, we complement the result of Proposition 4.4 by computing analytically the power of the test, which is the probability of rejecting *H* (no demographic change) when the data is produced under *H*^*′*^ (one change in the recent past). Recall that the test we use is a likelihood-ratio test with confidence level *α* = 0.95. It rejects *H* if 2Δlog ℒ > *z*_*α*_, where *z*_*α*_ is the *α*-quantile of a *χ*^2^ random variable with 2 degrees of freedom. In particular, *z*_*α*_ = 5.99 for *α* = 0.95.

Recall from Eq (20) that *q*_*w*_ is the *w*-quantile of the standard normal distribution.

#### Proposition 4.5

(Type II error). *The power P*_*α*_ *of the test, which is the probability of rejecting H under H*^*′*^ *by inferring rightly a deficit or excess of singletons as caused by a recent decline or expansion in population size, is larger than w iff the number k of generations since the change is*

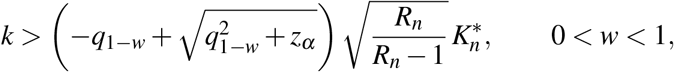

*where* 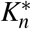 *was given in Equation* (19) *and R*_*n*_ *is the partial sum of the harmonic series* 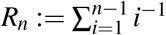.

#### Remark 4.6.

*Each of the previous two propositions ensures that the theoretical limit to the detection of a very recent demographic change is a minimal number k of generations which is equal to* 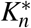 *times some bounded, positive statistical coefficient*.

*Proposition 4.4 states that the SFS realized under H is statistically different from the expected SFS under H*^*′*^ *with confidence level* > *w (type I error with probability* < 1 −*w*), *if* 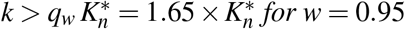 *for w* = 0.95.

*According to Proposition 4.5, the likelihood-ratio test detects (with confidence level α) that the SFS realized under H*^*′*^ *is different from the expected SFS under H, with probability* > *w (type II error with probability* < 1 −*w), if* 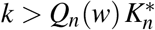, *where*

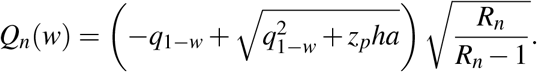

*Since* 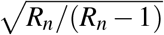 *decreases to 1 as n gets large (and is smaller than* 1.33 *as soon as n* ≥ 5*), we now approximate it by 1. Then for example, sticking to* 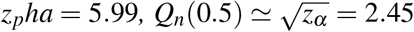 *(because q*_0.5_ = 0*) and Q*_*n*_(0.95) ≃ 3.25 *(because q*_0.05_ = −1.65).

*Note that when a type II error has a probability* 1 −*w close to 0 (power w close to 1), Q*_*n*_(*w*) ≃ −2*q*_1−*w*_ = 2*q*_*w*_, *which is surprisingly similar to the factor q*_*w*_ *obtained for a type I error to have probability* < 1 −*w*.

Before proving Proposition 4.5, we first need recalling some facts. We let (*f*_*i*_) and (*g*_*i*_) denote the normalized SFS under hypothesis *H* and *H*^*′*^ respectively.

The likelihood of a realized SFS (*ξ*_*i*_; 1 ≤ *i* ≤ *n* − 1) is defined in Equation (2), so that the log of the likelihood ratio is

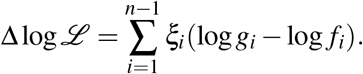

We assume that the realized SFS is obtained under the alternative scenario *H*^*′*^ with one (recent) demographic change corresponding to the expected normalized SFS (*g*_*i*_). In the setting of this work, this means that the number of segregating sites *S* is assumed given (and large), and that (*ξ*_*i*_; 1 ≤ *i* ≤ *n*− 1) has the same law as the multinomial random vector *<u>M</u>*(*S*) := (*M*_1_(*S*),…, *M*_*n*−1_(*S*)) with *S* trials and probabilities *<u>g</u>* = (*g*_1_, …, *g*_*n*−1_) to fall into bins 1, …, *n*− 1.

At the confidence level *α* (value used is *α* = 0.95), the likelihood-ratio test rejects *H* if 2Δlog ℒ > *z*_*α*_.

The power *P*_*α*_ of the test is then

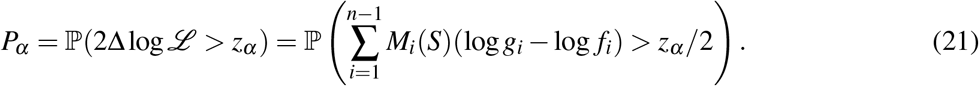

We first state a lemma leveraging central limit theorems for multinomial distributions.

#### Lemma 4.7.

*If for each i, there is b*_*i*_ *such that*

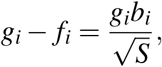

*then* 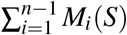 (log *g*_*i*_ − log *f*_*i*_) *converges as S* → ∞ *to the Gaussian random variable with mean β/*2 *and variance β, where*

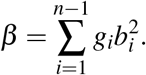

*Proof of Lemma 4.7*. It is well-known that 𝔼 (*M*_*i*_(*S*)) = *Sg*_*i*_ and Cov(*M*_*i*_(*S*), *M*_*j*_(*S*)) = *SC*_*ij*_, where

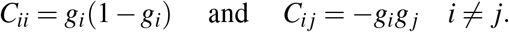

Also the central limit theorem ensures that 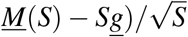 converges in law to the centered Gaussian vector with covariance matrix *C*. Then for any vector *a* = (*a*_1_, …, *a*_*n*−1_), the random variable

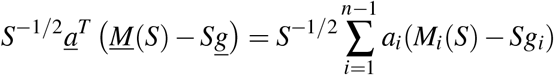

converges to the centered Gaussian variable with variance *β* given by

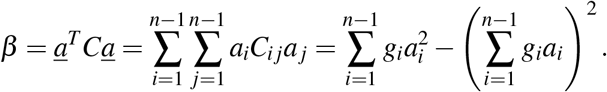

Now recall that for each *i*, we have 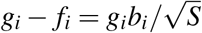, so that in particular

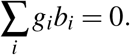

Then

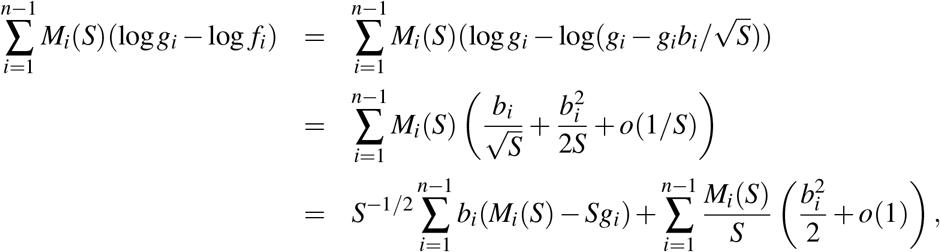

which, replacing *<u>a</u>* with *<u>b</u>* in the preceding central limit theorem, converges in law to the Gaussian random variable with mean

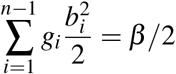

and variance

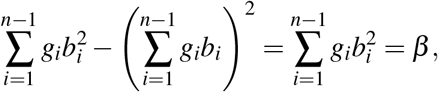

which ends the proof of the lemma.

We can now prove Proposition 4.5.

*Proof of Proposition 4.5*. We define (*x*_*i*_) and (*y*_*i*_) the expected SFS under hypothesis *H* and *H*^*′*^ respectively,

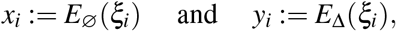

so that

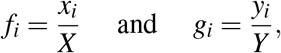

Where

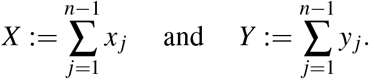

We seek to apply Lemma 4.7 to the case when the demographic change is recent. We always have *X, Y* and *S* of the same order and we will assume that 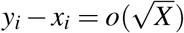 for each *i* ≥ 2 and

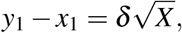

where *δ* can be positive (excess of singletons) or negative (deficit of singletons). Then 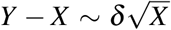, *f*_1_ = *x*_1_*/X* and

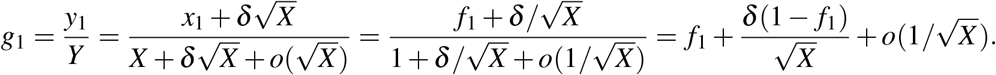

In addition, for any *i* ≥ 2,

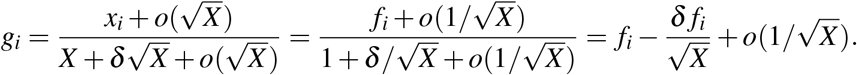

Now we seek *b*_*i*_ such that 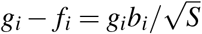, which, because 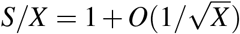, leads to

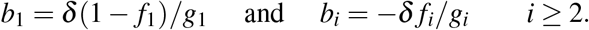

Note that we indeed recover ∑_*i*_ *g*_*i*_*b*_*i*_ = 0 and

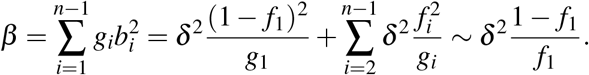

Now recall from Equation (21) that 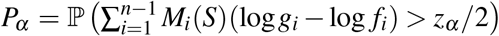 (log *g*_*i*_ − log *f*_*i*_) > *z*_*α*_ */*2, which thanks to Lemma 4.7 converges when the number of mutations/length of the sequence is large, to

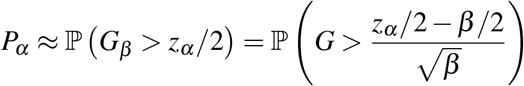

where *G*_*β*_ is a normal random variable with mean *β/*2 and variance *β* and *G* a standard normal variable (mean 0 and variance 1). Denoting by *q*_*w*_ the *w*-quantile of *G* (i.e. such that *P*(*G* < *q*_*w*_) = *w*), we get

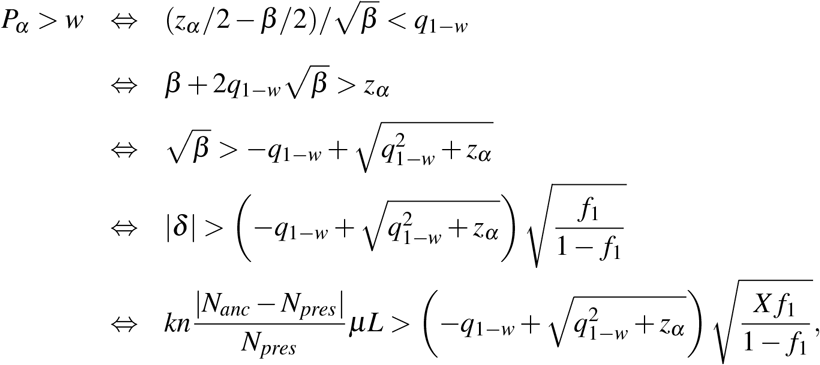

where the last line is due to Eq (13). Now with 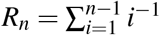, it is well-known that *f*_1_ = 1*/R*_*n*_ and because the total expected length (in number of generations) of the tree under *H* is 4*N*_*anc*_*R*_*n*_, we also have

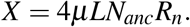

Then we get

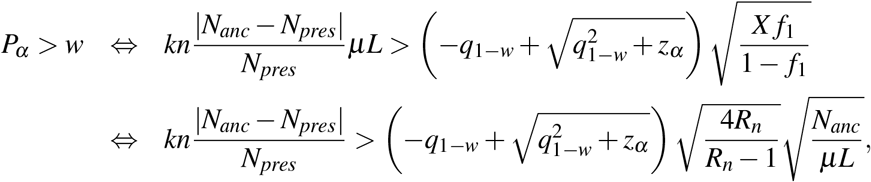

which ends the proof.

Proposition 4.4 stated that if a demographic change occurred 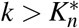 generations ago, then the first bin of the SFS is sufficiently distorted for the change to be statistically detectable. Now we claim that the second bin must also be sufficiently distorted, for the change to be statistically identifiable and correctly inferred. The next proposition ensures that this holds when 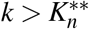, where

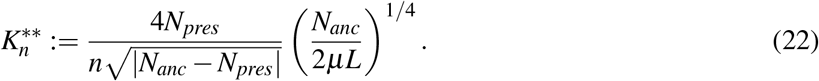

#### Remark 4.8.

*Assuming for example that N*_*pres*_ = *N*_*anc*_*/*2 *(population halving) and µL* = 2 *(as in Remark 4.3), then this second threshold number of generations is* 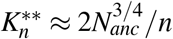.

#### Proposition 4.9

(2nd-order type I error). *The probability of attributing a deficit (resp. excess, resp. deviation) of doubletons to a false population decline (resp. expansion, resp. change), k generations ago, is larger than w if*

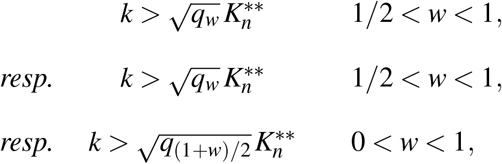

*where q*_*w*_ *is the standard normal w-quantile defined in Eq (20)*.

*Proof*. Similarly as in the proof of Proposition 4.4, under hypothesis *H*, 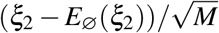 behaves like a centered normal random variable with variance 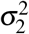 given by^1^

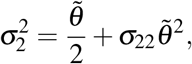

where, as soon as *n* > 4,

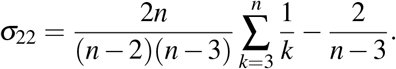

Here again, we can pick *ℓ* sufficiently small that 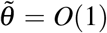 and so 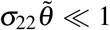, since *σ*_22_ only depends on *n* and goes to 0 as *n* → ∞. Then we have 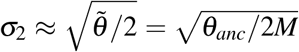. And again, assuming a population decline Δ time units ago,

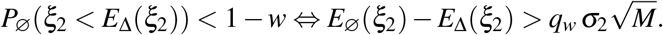

Replacing the two sides of the last inequality with their expressions yields

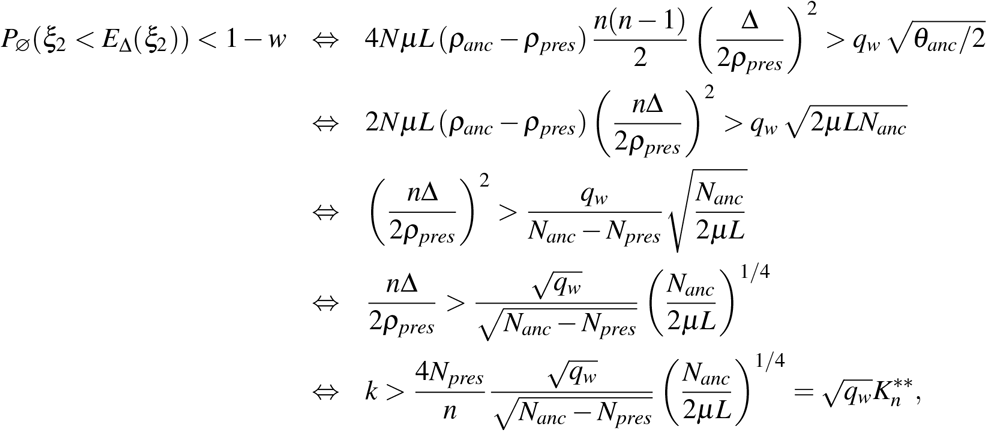

where *k* = 2*N*Δ.

### 5 Fitting complex scenarios of demography

In this section, we ask whether a simple strategy can improve the inference of complex demographic scenarios, such as the well-known zigzag model, i.e., piecewise-exponential.

First, three different SFSs with sample sizes of 100, 200, and 500 were simulated using the *stdpopsim* library^20,21^. Inference then proceeded as follows: the classical maximum-likelihood estimator proposed in *Blockbuster* was applied to the observed SFS to select the number of epochs. In a second step, inspired by the benchmark performed in the main text, 400 new SFS were generated by resampling the observed SFS using a Poisson distribution, in which each bin is centered at the observed value with variance equal to the mean. For each replicate, the maximum-likelihood step was performed using the number of epochs selected from the observed SFS. To keep computational time reasonable in this setting, the initial *Blockbuster* time grid size was fixed at *H* = 20 and the maximum number of epochs was set to *Q* = 7. The colored lines in Figure S1 represent the harmonic mean of the curves obtained across these replicates. According to the classical *Blockbuster* output, the number of epochs is 5, 7, and 6 in the inferred piecewise-constant model for sample sizes of 100, 200, and 500 haplotypes, respectively.

The number of changes in the trend direction is correctly inferred for the two larger sample sizes. For the smallest sample, the direction is correctly recovered only if the 6-epoch model is used, although the estimated magnitudes and timing of changes may not be accurate. This result suggests that, in cases where prior knowledge of the underlying model exists, using more epochs than indicated by the likelihood-ratio test can still aid interpretation. In any case, this is consistent with previous findings that different models can yield similar SFS, especially when the sample size is small. The result for the SFS computed from 200 haplotypes is surprisingly closer to the true model than those from 500 haplotypes, for which the inferred curve is very similar to the piecewise-constant model. This can be explained by the smaller amount of uncertainty in the the 500-haplotype SFS, due to both the larger sample size and the smaller number of epochs (6 versus 7). We can conclude that such a sampling strategy may help fit continuous models, but the amount of noise introduced between replicates must be carefully adjusted: it should be increased when uncertainty is low, but not so much that it obscures the signal. Identifying an objective and principled method to achieve this is not trivial

**Figure S1:**
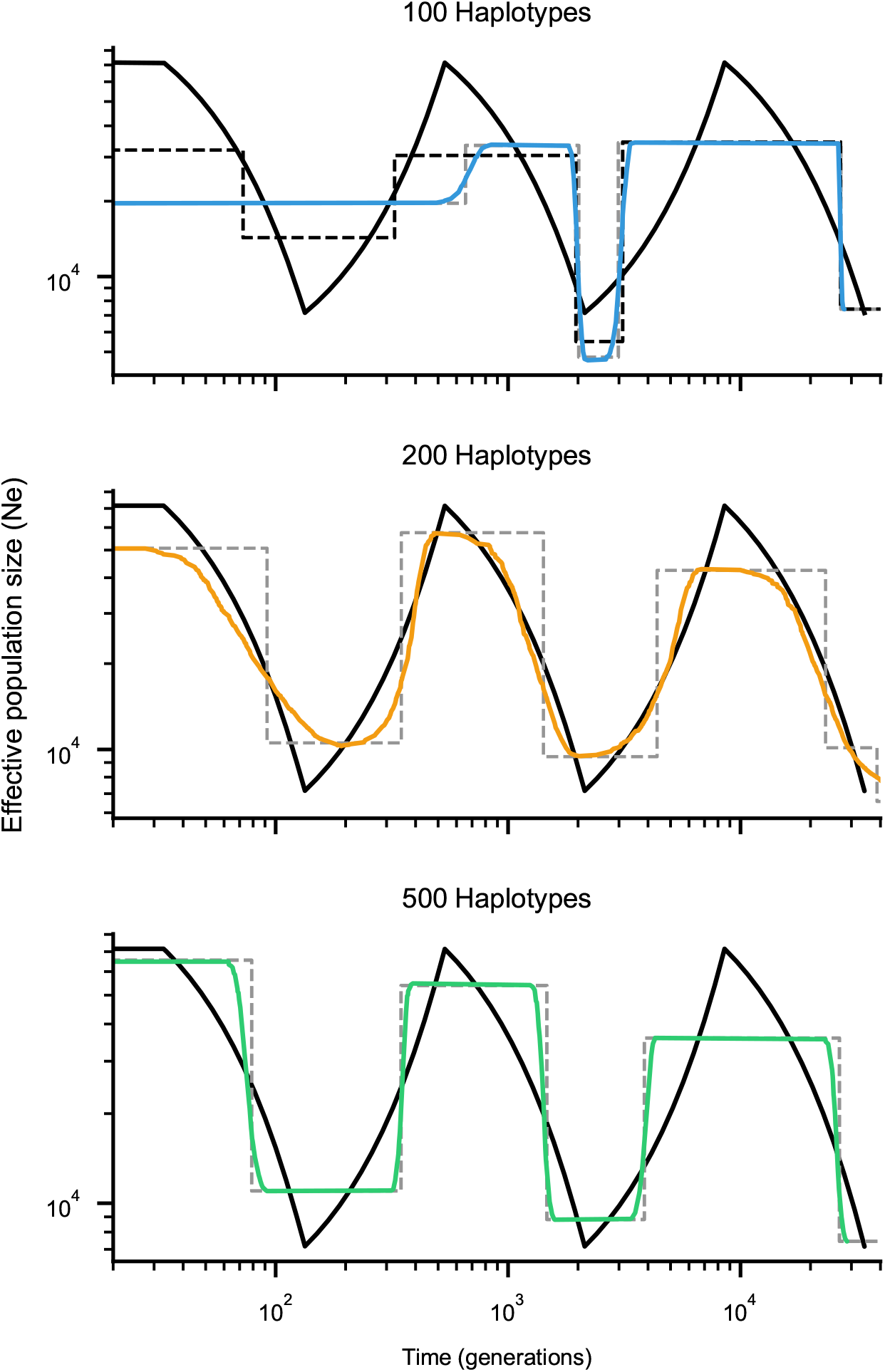
Inference over the zigzag model for three different sample sizes. Colored curves show the harmonic mean across 400 inferences on sampled SFS, and the dashed grey line shows the model inferred using the MLE estimation from *Blockbuster*.

